# Emergent phases of ecological diversity and dynamics mapped in microcosms

**DOI:** 10.1101/2021.10.28.466339

**Authors:** Jiliang Hu, Daniel R. Amor, Matthieu Barbier, Guy Bunin, Jeff Gore

## Abstract

Natural ecological communities display striking features, such as high biodiversity and a wide range of dynamics, that have been difficult to explain in a unified framework. Using experimental bacterial microcosms, we perform the first direct test of recent complex systems theory predicting that simple aggregate parameters dictate emergent behaviors of the community. As either the number of species or the strength of species interactions is increased, we show that microbial ecosystems transition between distinct qualitative dynamical phases in the predicted order, from a stable equilibrium where all species coexist, to partial coexistence, to emergence of persistent fluctuations in species abundance. Under the same conditions, high biodiversity and fluctuations allow and require each other. Our results demonstrate predictable emergent diversity and dynamics in ecological communities.

**One-Sentence Summary:** A phase diagram of ecological dynamics and diversity as a function of coarse-grained features of species interaction network.

---

Natural species reside and interact with myriad other species in complex communities (*1*). Central challenges of ecology include understanding what explains species diversity in communities, what determines their dynamical behaviors (*2–4*), and how diversity and dynamics interact to shape ecological services and functions (*5*, *6*). Field observations have uncovered a broad range of dynamical behaviors and their relationships with diversity in natural settings (*7*), which are often understood in the context of environmental drivers affecting both (*8*). Laboratory experiments in controlled environment offer the possibility of disentangling these external drivers from inherent community properties. Previous experiments have established the possibility of predictable dynamics (stable equilibria or limit cycle oscillations) in systems comprising a few species (*9–14*), which were explained by inter-species interactions such as predation (*12*, *13*, *15*), competition (*9*, *10*), and cross-feeding (*16*, *17*). Furthermore, there is evidence of more complex fluctuating dynamics at the species level, as well as predictable behaviors at coarse-grained taxonomic levels, in highly diverse experimental communities (*4*, *17*–*19*). Yet, it has been a challenge to discern general principles explaining the diversity and dynamics of complex multi-species communities, since detailed biological parameters are typically not available in large ecological networks. Therefore, a fundamental question remains: is it possible to identify simple aggregate parameters governing the macroscopic diversity and dynamics of ecological communities?

Starting with the pioneering work of Robert May (15), ecologists have sought to predict key emergent properties of complex communities based on coarse-grained features of the interaction network, such as the number of species and the distribution of interaction strengths between species. May and others have suggested that large or strongly interacting communities will generically be unstable (*20*–*26*), yet we still do not understand how such large communities may arise, nor the dynamical behavior of communities that are not stable. In particular, it has been shown that species can go extinct before the community loses stability (*27*–*30*), and also that unstable communities can display a range of dynamics including periodic (limit cycle) oscillations or chaotic fluctuations, that in some cases play a role in sustaining diversity (*31*–*37*). This body of theory has however been difficult to validate because driving parameters such as ecological interactions are often challenging to estimate (*38*, *39*). What has largely been missing is a controllable experimental setting where these theories can be tested systematically over a wide range of tunable conditions.

To guide our experiments, we begin by modeling the long-term dynamics and diversity of ecological communities using the well-known generalized Lotka-Volterra (gLV) model, modified to include dispersal from a species pool:

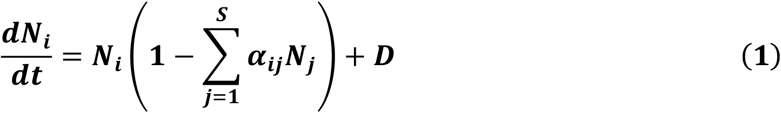

where *N_i_* is the abundance of species *i* (normalized to its carrying capacity), *α_ij_* is the interaction strength that captures how strongly species *j* inhibits the growth of species *i* (with self-regulation *α_ii_* = 1), and *D* is the dispersal rate from an outside species pool to the focal community. We simulated the dynamics of communities with different species pool sizes *S* and interaction matrices. We sample the interaction strength from a uniform distribution *U* [*0, 2<α_ij_>]*, where <*α_ij_*> is the mean interaction strength between species (we show later that our predictions are robust to different choices of distributions of interaction strengths, but depend on a combination of interaction mean and variance, here entirely controlled by parameter <*α_ij_*>). Modeling species interactions as a random interaction network captures the heterogeneity of species without assuming any particular community structure (*20*, *21*, *25*, *29*).

Our simulations revealed a strong dependence of the steady state diversity and dynamics of the community on both the species pool size *S* (Fig. 1A) and interaction strength <*α_ij_*> (Fig. 1B). As either of these parameters increase, communities experience a transition from I) stable full coexistence—all species survive and reach stable abundances, to II) stable partial coexistence—some species go extinct (Methods, Fig. S1), and the surviving ones reach stable abundances, to III) persistent fluctuations in species abundances. A linear stability analysis confirms that the transition to unstable dynamics (II → III) coincides with the community matrix eigenvalues exhibiting positive real parts (Fig. S2). Recent theory also proved analytically the existence of a phase transition from a unique stable state (I and II) to persistent fluctuations or alternative stable states (III) using the same control parameters (*27*). In accordance with these theoretical predictions (*27*, *28*, *40*), our simulations demonstrate that the size of the species pool *S* and the strength of interactions <*α_ij_*> combine to determine the dynamical behaviors exhibited by ecological communities.

**Fig.1.**
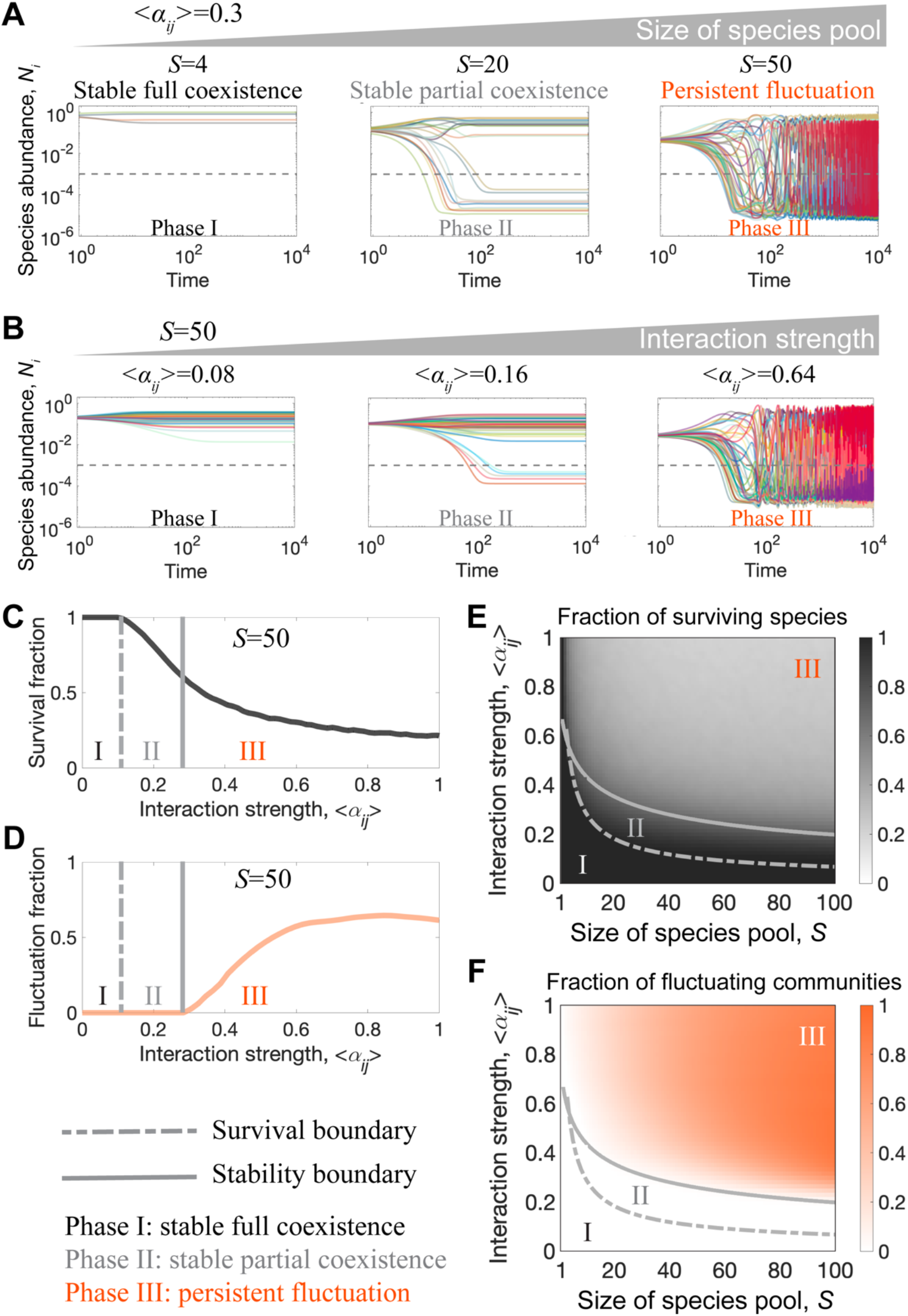
Theory predicts that species pool size and interspecies interaction strength shape phases of community diversity and dynamics. (A) Representative time series of species abundance for the qualitatively different dynamics of communities with different species pool size *S*, given the same interspecies interaction strength <*α_ij_*>=0.3. Communities transition from stable full coexistence (*S*=4) to stable partial coexistence (*S*=20) to persistent fluctuations (*S*=50). Analogous changes in community dynamics also result from increasing the interaction strength (B) while keeping the species pool size constant. (C) Mean fraction of species that survive in the community, and fraction of communities that exhibit persistent fluctuations (D) as a function of the interaction strength. As interaction strength increases, communities lose species (transition from phase I to II, vertical dashed line) before losing stability (transition from phase II to III, solid vertical line). Mapping the survival fraction (E) and community fluctuation fraction (F) onto the phase space reveals that the sequence (phase I to phase II to phase III) of phase transitions is maintained as either of the control parameters increases. The gray dashed (solid) line shows the analytical solution for the survival (stability) boundary, and the color maps account for numerical results (Supp. Material).

To further understand the ecological implications of the different dynamical phases in our model, we analyzed both the fraction of species that survive in the long term (Fig. 1C, E) and the fraction of communities that exhibit persistent fluctuations (Fig. 1D, F) over a wide range of parameter values. This analysis confirmed that the observed sequence of three dynamical phases is a generic trend across the parameter space: communities generally experience species extinctions before they lose stability (Fig. 1C-F) as either the species pool size or the interaction strength increase. Importantly, this order of transitions between dynamical phases—stable full coexistence, stable partial coexistence, and persistent fluctuations—is not only predicted by analytical expressions for the phase boundaries (Fig. 1C-F), but it is also robust to different choices of interaction strength distributions and modeling assumptions (Fig. S3) (*27*). Our results suggest that it may be possible that the diversity and dynamics of multi-species communities to be predicted by coarse-grained features—the statistics of interaction strengths and species pool size—without detailed knowledge of the interaction network or the underlying mechanisms behind those interactions.

To experimentally test the theoretically predicted transitions between dynamical phases in ecological communities, we built synthetic microbial communities using a library of 48 bacterial isolates from terrestrial environments (Materials, Fig. S4, 5). This library is phylogenetically diverse, with isolates coming from 26 different families among 4 phylums: Proteobacteria, Firmicutes, Bacteroidota and Actinobacteriota. We used subsets of this library to obtain species pools of different sizes, which we exposed to serial cycles of growth and dilution in the presence of dispersal from the species pool (Fig. 2A). To monitor the dynamics of these communities, we measured community composition via 16S ribosomal RNA (rRNA) amplicon sequencing and total biomass via optical density (OD) at the end of each daily cycle. In addition to varying the size of the species pool, we experimentally tuned the strength of the interactions between community members by varying the nutrient concentration in the media. Consistent with previous work (*19*, *41*), we found that the probability of coexistence in pair-wise co-culture decreased as we supplemented the media with glucose and urea (Fig 2B). In this media, an increase in nutrient concentration therefore increases the strength of competitive interactions. This experimental platform allows us to control the two key parameters established by theory—the species pool size and the interaction strength between species—to drive communities between different dynamical phases.

**Fig.2.**
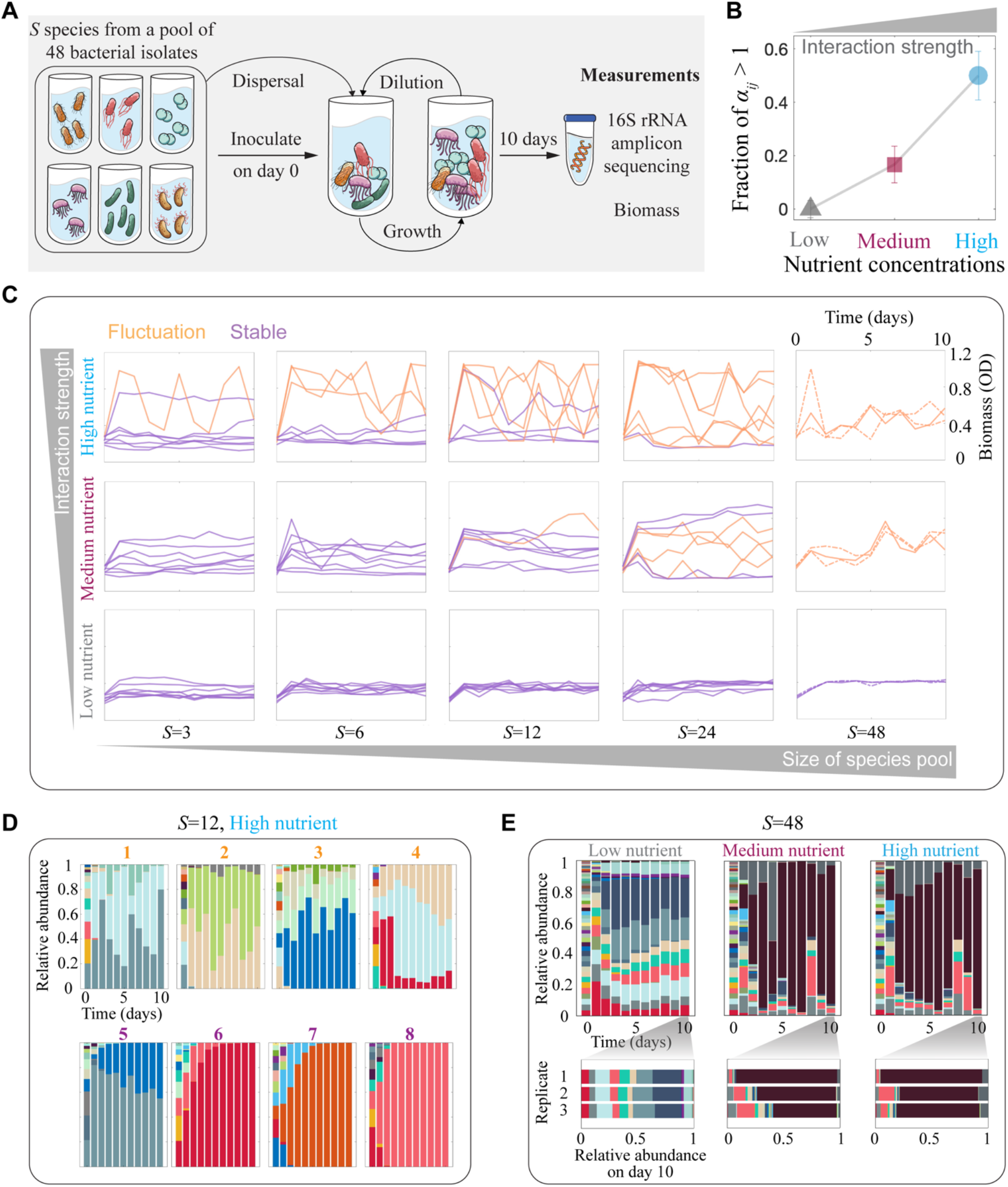
Increasing species pool size or interaction strength leads to loss of stability in experimental microbial communities. (A) We constructed subsets of a library of 48 bacterial isolates to create experimental species pools of different sizes and compositions. We used these species pools to inoculate communities that were exposed to 10 daily cycles of growth and dilution with dispersal from the species pool. To monitor community dynamics, we measured community composition (via 16S sequencing) and total biomass (through optical density, OD) at the end of each cycle. (B) Measuring the experimental outcome of pairwise (2-species) competitions revealed that the fraction of competitive exclusion (one of the competitor species goes extinct) cases, and, therefore, the fraction of interaction strengths that leads to the loss of pairwise coexistence (*α_ij_* > 1), increased with nutrients concentration. This result enables using different nutrient (glucose and urea) concentrations to experimentally tune the interaction strength (see Supp Material). Error bars, s.e.m., n=30. (C) Fluctuations in microbial community biomass increase with either species pool size or interaction strength. Solid lines stand for 8 different species pool compositions, and dashed lines in the *S*=48 case show the time series for 2 additional replicates with identical species pool composition. Purple (orange) lines highlight stable (fluctuating) dynamics between days 7 and 10 (see Supp Material). (D) 16S sequencing data reveals that, under strong interactions (high nutrients concentration), half of the 12-species communities exhibit persistent fluctuations (top panels) in species abundances, and the other half reached stability (bottom panels). Column colors represent specific amplicon sequence variants (ASV), which correspond to different species of the library. (E) Time series (top panels) for the species abundances in a 48-species community in the different interaction strength (nutrients concentration) conditions. The 48-species community reached stability only in the low interaction strength (low nutrients concentration) condition. Community composition at the end of the experiment was also highly reproducible in this condition, while higher interaction strengths led to higher variability between replicates (bottom panels, and Fig. S13-15).

We experimentally mapped the phase space of community dynamics by exposing three replicates of 189 synthetic communities of different species pool sizes (*S* = 2 – 48) to three levels of interaction strength. The time series for the total biomass of these communities were relatively stable when the interaction strength was low and species pool was less diverse, while increasing these two variables progressively led to a higher fraction of communities exhibiting biomass fluctuations (Fig. 2C). Analyzing species abundances through 16S sequencing, we found that total biomass fluctuations were highly correlated with species abundance fluctuations in these synthetic communities (Fig. S6). For example, for communities with 12 species in the pool and high nutrient concentration, 4 out of 8 communities reached stable equilibria, and the remaining 4 exhibited fluctuations in both biomass and species abundances until the end of the experiment (Fig. 2C and D). We performed three replicates for each community we studied here, and we found that the replicates of the same communities exhibited highly reproducible dynamics (Fig. S7-15). The classification of communities into fluctuating and stable groups was robust to different methods based on biomass or species compositions, and agreed with a classification through divergence between replicates. (Methods, Fig. S6). Single species monoculture (*S*=1) and pairwise co-culture (*S*=2) always reached stable equilibria in our experiment (Fig. S16). These results show that our synthetic microbial communities lose stability as either the species pool size or the interaction strength is increased, as predicted by theory (Fig. 1).

To understand how this loss of stability is related to species extinction, we analyzed the fraction of species surviving in the different experimental conditions. As expected, we observed a decrease in the fraction of species that survived as we increased either the species pool size or the interaction strength—determined by nutrient concentration (Fig. 3A). For example, at medium interaction strength 83% (+/- 3%) of species were able to survive in the fifteen pairwise co-cultures tested (*S* = 2), and this survival fraction decreased to 36% (+/- 7%) among the eight different combinations of six species communities (S = 6) (Fig. 3A). Despite the significant loss of species, none of these communities displayed persistent fluctuations (Fig. 3B). Only with further increase of the species pool size did we begin to observe fluctuations in species abundance, with half of the 24-species combinations displaying fluctuations (Fig. 3B). Interestingly, the species survival fraction displayed only a modest decrease entering the fluctuation regime, with 24% (+/- 2%) of species surviving in the 24-species communities down from 36% (+/- 7%) surviving in the 6-species communities (Fig. 3A and B). Mapping these experimental results over the phase space (Fig. 3C and D) confirmed that transitions between dynamical phases occur in the specific order predicted by theory (Fig. 1E and F): communities experience species extinctions before exhibiting persistent fluctuations, as either species pool size or interaction strength increases.

**Fig.3.**
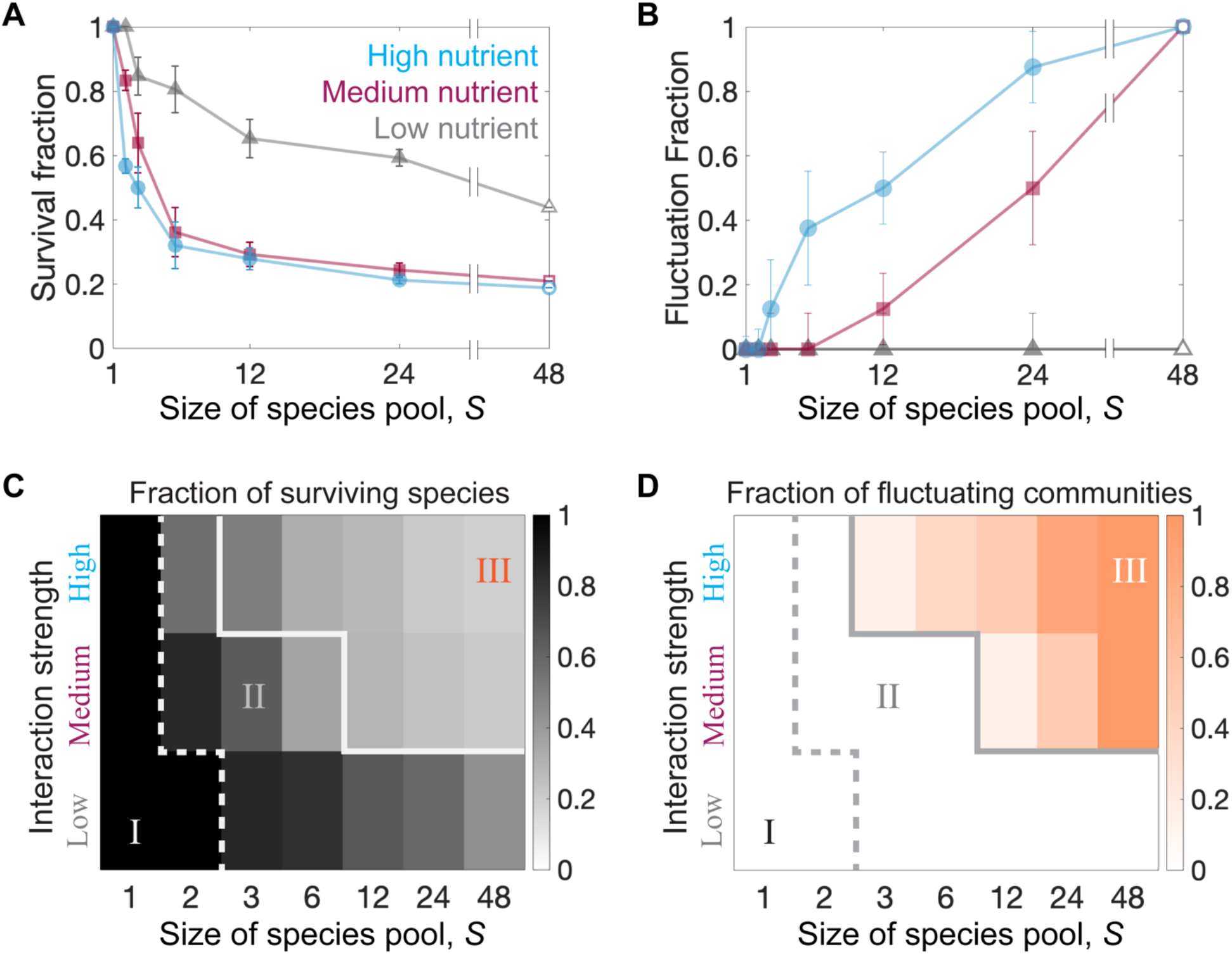
Species pool size and interaction strength determines the diversity and dynamics of experimental communities. (A) The fraction of surviving species decreases as either the species pool size or the interaction strength—determined by nutrient concentration—increase. The survival fraction decreases more slowly at high *S* and strong interaction strength. Error bars, s.e.m., n=8. (B) The fraction of fluctuating communities increases as we increase either the species pool size or the interaction strength. Error bars, s.e.m., n=8. (C) Phase diagram for the fraction of species surviving in experimental communities. The dashed line shows boundary of phase I, where all species survive. As communities move farther away from phase I, they experience species extinctions, with a fast decay in survival fraction through phase II, and a relative maintenance of survival fraction through phase III. (D) Phase diagram for the fraction of fluctuating communities in experiments. The solid gray line indicates the boundary between phase II and phase III, where communities start exhibiting persistent fluctuations. Experimental communities universally exhibit the three predicted phases of stable full coexistence (I), stable partial coexistence (II) or persistent fluctuations (III) as a function of both species pool size and the interaction strength.

To address the question of how fluctuations and diversity affect each other, we analyzed the species survival fraction reached by individual communities with different species compositions in the same conditions. In simulations, the fraction of surviving species revealed a generic trend: for the same species pool size and interaction strength, fluctuating communities were more diverse than stable communities (Fig. 4A). This trend was also observed in experimental communities, where the vast majority of fluctuating communities reached higher survival fractions than stable communities under the same experimental conditions (Fig 4B). For example, in the 12-species communities, the fluctuating communities had on average 5 +/- 1 species surviving, as compared to only 2+/- 1 species surviving in the stable communities. Among the fluctuating communities, 88% (+/-5%) exhibited survival fractions above or equal to the mean, as compared to only 14% (+/- 6%) in the case of stable communities (p<0.05, Methods). Both experiments and simulations suggested that fluctuations are an emergent, diversity-dependent phenomenon, as we selected pairs of stable communities, combined their species pools to assemble composite communities, and found that some exhibited fluctuations (Fig. S17). We also found numerically that fluctuations and high diversity disappeared together as we stopped the dispersal or pinned the abundance of the most abundant species (Fig. S1). Our results suggest that persistent fluctuations and high diversity require and allow each other, as theoretically shown in previous work (*32*, *33*).

**Fig.4.**
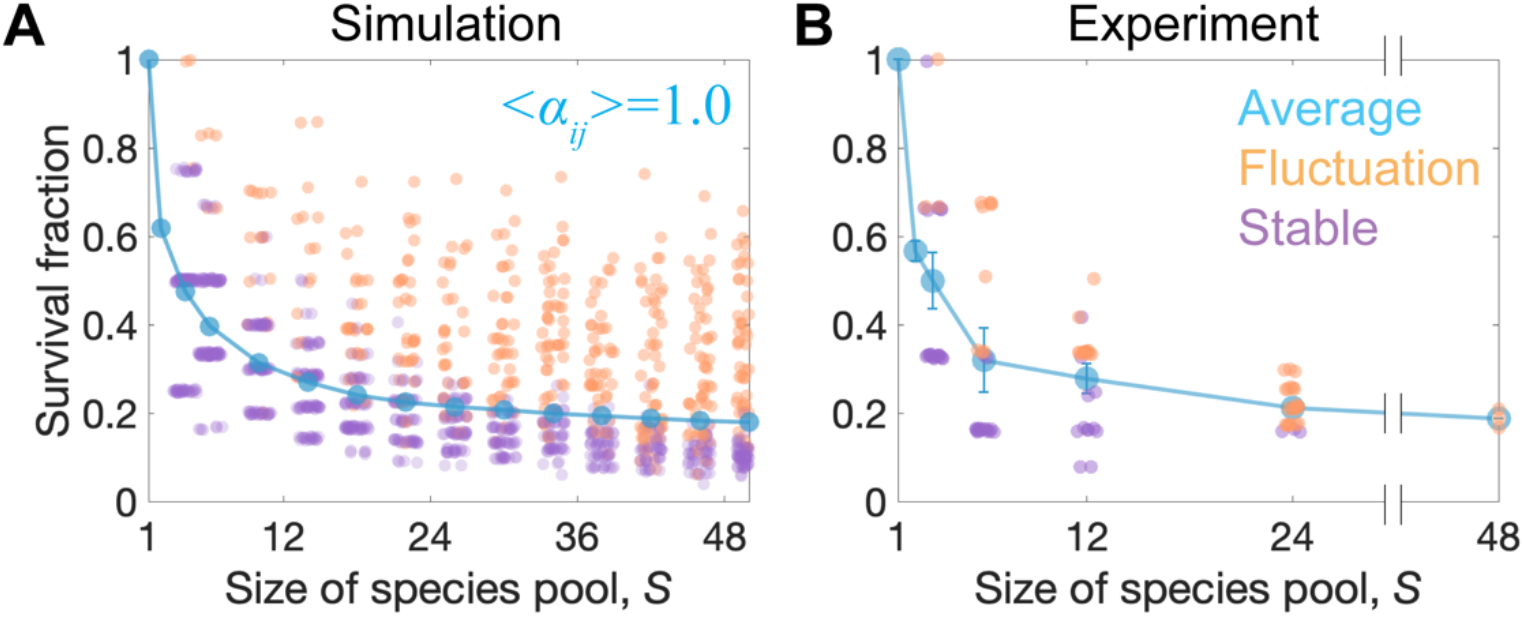
Fluctuating communities are more diverse than stable communities under the same conditions. (A) As the average survival fraction decreases with increasing species pool size S in simulations, more communities exhibit fluctuations in species abundances (orange points). While stable communities (purple) exhibit a steady decrease in species survival fraction as the species pool size increases, the loss of species is slower in fluctuating communities. Each point represents an individual community. (B) In experiments, fluctuating microbial communities also exhibit a higher survival fraction than stable communities. The survival fractions of 88% (+/-5%) of the fluctuating communities are above or equal to the mean, as compared to 14% (+/- 6%) in the case of stable communities (p<0.05, Methods); error bars, s.e.m., n=8.

Our study contributes insight into the long-debated relationship between diversity and stability. We found a strong positive correlation between realized diversity (number of surviving species) and instability (abundance fluctuations). This finding is consistent with two major ideas in theoretical ecology: in one direction, May’s suggestion that complexity leads to instability (*20*), and in the other direction, Chesson’s argument that temporal fluctuations can help maintain diversity (*42*). Our results suggest that both of these mechanisms are at play, with both diversity and instability promoting each other, which is consistent with recent theory (*33*).

We uncovered this diversity-stability relationship by experimentally controlling two factors that are usually unobservable in natural settings: the pool size of species that may invade a community, and the interaction strength (here tuned via resource levels). Both factors could play a confounding role by varying across ecosystems, affecting diversity and stability simultaneously. Variation in interaction strength could even reverse the realized diversity-stability relationship. On the one hand, stability decreases with increasing either size of species pool or realized diversity (number of surviving species) for any given value of the interaction strength (Fig 1F, 3D, 4A and 4B). On the other hand, increasing interaction strength for a given species pool size leads to a lower realized diversity and lower stability fraction (Fig. 1E and 3C), creating a positive correlation between diversity and stability. We believe that these different interplays between parameters across orthogonal directions in the phase diagram could underlie some of the seemingly contradictory results from field experiments (*8*) addressing the diversity-stability relationship.

Furthermore, our experimental setting allowed us to isolate the community dynamics from environmental fluctuations. While many ecological communities exhibit abundance fluctuations, the question of whether such fluctuations are inherent to the community—arising from species interactions—or are instead driven by external factors has seldom been addressed empirically and systematically. Under laboratory conditions that minimize environmental fluctuations, our results suggest that two inherent components of an ecological community—the size of species pool and the strength of interactions—can determine both the stability and long-term biodiversity reached by an ecosystem. Our experimental results agree with predictions of recent theory (*27*, *29*, *43*) that coarse-grained features of the interaction network are sufficient to predict the dynamical behaviors of complex ecological communities. These predictions are robust to various biological ingredients (e.g., intraspecific diversity and inter-species interaction mechanism), and can also be recapitulated in a resource-explicit model (*44*). Therefore, the emergent phases of diversity and dynamics that we observed here may occur in a wide range of ecological communities. Future work should strive to determine whether these emergent phases generalize across spatiotemporal scales, environmental conditions and organism types, to understand their prevalence and importance in shaping major ecological patterns (*45*).

## Acknowledgments

We thank all the Gore Lab members for inspiring discussions. We thank Martina Dal Bello at MIT for drawing the cartoons of bacterial cells. We also thank Christoph Ratzke, currently at University of Tübingen, for the isolation of bacterial species, as well as the Eric Alm laboratory at MIT for sharing laboratory equipment. We thank the groups led by Daniel Fisher at Stanford and Oskar Hallatschek at UC Berkeley for valuable discussions.

## Author contributions

J.H. and J.G. conceived the study. J.H. and D.R.A. performed the experiments. J.H., M.B. and G.B. performed the theoretical modelling. All authors analyzed the data and wrote the manuscript.

## Data and materials availability

Isolates and communities are available upon request. All data are available in the main text or the supplementary materials.

## Supplementary Materials

Materials and Methods

Figs. S1 to S18

References (*46*–*50*)

