## Supplementary Materials for "Emergent phases of ecological diversity and dynamics mapped in microcosms"

**This PDF file includes:**

Materials and Methods
Figs. S1 to S18

### Materials and Methods

Bacterial isolates, media and culturing conditions

We constructed the library of 48 bacterial species using 24 bacterial isolates from soil samples taken at Middlesex Fells Reservation in Somerville, Massachusetts, and 24 isolates from the *C. elegans* intestine (*46*).

In the case of low interaction strength (low nutrients concentration) conditions, experimental communities were cultured in Base Medium (BM): 1gL$^{-1}$ yeast extract and 1 gL$^{-1}$ soytone from Becton Dickinson, 10 mM sodium phosphate, 0.1 mM CaCl2, 2 mM $MgCl_2$, 4mgL$^{-1}$ $NiSO_4$ and 50 mgL$^{-1}$ $MnCl_2$, pH adjusted to 6.5. For intermediate interaction strength (medium nutrients concentration) conditions, we used BM supplemented with 5 gL$^{-1}$ glucose and 4 gL$^{-1}$ urea. For the high interaction strength (high nutrients concentration) condition, we used BM supplemented with 20 gL$^{-1}$ glucose and 16 gL$^{-1}$ urea. All media were filter sterilized using Bottle Top Filtration Units (VWR). All of the chemicals were purchased from Sigma–Aldrich unless otherwise stated.

Both monocultures and communities of the bacterial isolates were grown in 96-deepwell plates (Deepwell plate 96/1000µl; Eppendorf) covered with AeraSeal adhesive sealing films (Excel Scientific). The incubation temperature was 30 °C for all communities. The deepwell plates were shaken at 1,200 r.p.m. on Titramax shakers (Heidolph). To minimize evaporation, the plates were incubated inside custom-built acrylic boxes.

Pre-cultures, daily dilutions, dispersal, and biomass measurements

Before each experiment, pre-cultures were initiated by thawing the bacteria and inoculating individual species into 600 $\mu$L of BM. The resulting monocultures were exposed to 5 daily cycles of growth and (30-fold) dilution into fresh media. At the beginning of each experiment, aliquots of these monocultures were mixed in equal volume proportions to form the synthetic communities. During the experiment, the monocultures were exposed to further dilution cycles and used to apply the daily dispersal into the synthetic communities as described below.

We created 189 different synthetic communities using randomly generated subsets of the library of isolates, each subset constituting the species pool (of size *S*) for each community. After mixing monocultures in equal volumes, each experimental community was initiated by inoculating 20 $\mu$L of its initial mix of isolates into 600 $\mu$L of BM, repeating the process to generate a total of 3 replicates per community. To form the communities with S≤12, we created subsets of random species from the soil isolates, and for S>12 we randomly matched both *soil* and *C. elegans* isolates. The resulting synthetic communities were cultured under serial dilution cycles with dispersal as follows.

To apply a $10^{-6}$ dispersal rate, every 24hr monoculture aliquots of the species in each community pool were mixed at equal volumes, and then diluted by a $10^4$ factor before inoculating 6uL of this mix into the wells containing the corresponding experimental community matching each species pool. After this, the experimental cultures were thoroughly mixed using a 96-well pipettor (Viaflo 96, Integra Biosciences; settings: pipette/mix program, 5 mixing cycles, mixing volume 300 µL, speed 6) before applying a 30-fold dilution by transferring 20 μL of the cultures into a new plate with 600 μL of fresh media.

Experiments were extended to a total of 10 daily cycles. At the end of every daily cycle, 150uL samples of each culture were used to measure the OD (600nm), a proxy for the total biomass in the cultures, using a Varioskan Flash (Thermo Fisher Scientific) plate reader. The remaining culture volume was stored at -80 °C for subsequent DNA extraction.

DNA extraction, 16S rRNA sequencing and data analysis.

DNA extraction was performed with the QIAGEN DNeasy PowerSoil HTP 96 Kit following the protocol provided by the manufacturer. The obtained DNA was used for 16S (V4 region) amplicon sequencing. Library preparation and Illumina MiSeq sequencing were performed by the Environmental Sample Preparation and Sequencing Facility at Argonne National Laboratory. We used the R package DADA2 to obtain the amplicon sequence variants (ASVs) as described by Callahan *et al*.(*47*). Taxonomic identities were assigned to the ASVs by using SILVA (version 132) as a reference database. For each sample. species richness was calculated as the number of ASVs with a relative abundance $\geqslant 0.1\%$, which corresponds to the $0.1\%$ extinction threshold used in simulation (Fig. S1). The phylogenetic tree (Fig. S5) was constructed using Simple Phylogeny (*48*) by the EMBL's European Bioinformatics Institute. Taxonomic identities were assigned to ASVs using Randomized Axelerated Maximum Likelihood (RAxML) using default parameters.

Numerical methods

All simulations used the Runge-Kutta method on Matlab to numerically solve the LV equations (with an integration step of 0.05). A definition of 100 x100 pixels was used for each phase diagram, linearly segmenting the parameter space in the ranges $<\alpha_{ij}> \in [0, 1.5]$ and $S \in [1, 100]$. In each phase diagram, each pixel shows the average result for $10^3$ simulations. The total simulation time is $10^4$.

Extinction threshold *in silico*

The presence of the dispersal rate in Eq. (1) makes species to exhibit strictly positive abundances in Fig 1B and C. Nevertheless, we consider that a species is extinct if its abundance lays below a $10^{-3}$ threshold. Around this threshold, the dispersal rate becomes the main effector preventing abundance decay (Fig. S1).

Definition of stable and fluctuating dynamics *in silico*

To differentiate between stable and fluctuating communities, we computed the average coefficient of variation of $N_i$ between $t=5\times10^3$ and $t=10^4$. We define communities with this average coefficient of variation higher (lower) than $10^{-3}$ as fluctuating (stable) communities (Fig. S1).

Survival fraction *in silico*

To compute the survival fraction, we computed the fraction of species whose abundance exceeded the extinction threshold at any time during the last 100 units of time in the simulation. Our choice of including a time window when measuring diversity is motivated by the fact that, for the case of unstable communities, species abundances fluctuate above and below the extinction threshold over time. Since we measured diversity and species compositions every 24 hours in the experiment, we consider an analogous window of 100 time units in simulations.

Analytical curves for boundaries between phases, and sharpness of the transitions

The analytical boundary between the stable phase (II) and the fluctuating phase (III) was derived in Bunin 2017 (*27*). For equal carrying capacities, it is shown that the boundary lies at the average standing species richness $S^* = S/2$, when $\sigma \equiv \sqrt{S}\mathrm{std}(\alpha_{ij})/(1 - \langle\alpha_{ij}\rangle) = \sqrt{2}$. There and in (*49*) it is also shown that the loss of stability of the equilibrium coincides with real parts of some community matrix $(-\alpha_{ij})$ eigenvalues becoming positive. For any distribution of interaction strengths (uniform, exponential, *etc.*), $\mathrm{std}(\alpha_{ij})$ and $\langle\alpha_{ij}\rangle$ can be calculated, and this criterion applied. The boundary between the fully-coexisting (I) and stable (II) phases is given by taking the prediction for the average standing species richness $S^*$ given in (*27*), and setting $S^* = S - 1$, namely the parameters when one species has gone extinct. This is not expected to be exact at large $S$, since the prediction in (*27*) is exact in conditions where $S^*/S$ is finite at large $S$, while here the boundary is at $S^*/S = 1 - 1/S$ which approaches 1 at large $S$. Nonetheless, it gives good results in the range of pool diversities shown in Fig. 1. Other techniques for analyzing the transition are also possible (*50*).

*Sharpness of transitions* – The term "phase transition" in physics implies that the transition is sharp in systems with many degrees of freedom. The transition between phases (II) and (III) is known to be sharp (*27*, *28*): at high $S$, communities that lie above the phase boundary (e.g., $\sigma > \sqrt{2}$ with all species exhibiting equal carrying capacities) always (with probability one) exhibit persistent fluctuations, while communities below the phase boundary (e.g., when $\sigma < \sqrt{2}$ for equal carrying capacities) reach a stable equilibrium .

The transition between phases (I) and (II) is also sharp when $S$ is large, in the following sense. Fig. S18 shows the probability of full coexistence as a function of $\langle\alpha_{ij}\rangle$, for different values of $S$. The x-axis is normalized by $\langle\alpha_{ij}\rangle$ where the analytical boundary is expected. This makes all curves decrease to zero in the same region, but the width of the crossover regime becomes narrower with increasing $S$. In other words, the width of the crossover region between the phases is small compared to the width of phase (I), for large $S$.

The fact that all curves decrease to zero at the same region, shows that the analytical expression indeed captures the correct dependence of the boundary in $\langle\alpha_{ij}\rangle$ on $S$. The fact that the crossover happens around a value of ~2.1 rather than around 1, is due to the inexact theory used, as explained above.

Definition of stable and fluctuating experimental communities

To differentiate between stable and fluctuating communities in experiments, we computed the average coefficient of variation for species abundances from day 7 to day 10. This corresponds to the average value of the standard deviation for the abundance of each species $N_i$ (over day 7, day 8, day 9 and day 10) scaled by average species abundance. Communities for which the average coefficient of variation in this time range is below (above) a 0.25 threshold are considered stable (fluctuating) communities (Fig S6). We also find that the biomass and species compositions fluctuate asynchronously across replicates of fluctuating communities starting from the same initial conditions, thus a large difference of compositions across different replicates under the same condition can be another indicator of community fluctuations. The cluster of communities' dynamics is consistent between the metric by average coefficient of variation and by final composition difference across replicates, as shown in Fig. S6.

Quantification and statistical test of species survival fractions

To statistically test differences in species survival fractions between fluctuating communities and stable communities in Fig. 4B, we first calculated the difference of survival fraction between each community (purple or orange points in Fig. 4B) and the corresponding average survival fraction (blue points in Fig. 4B) for each species pool size (S). We then performed an analysis of variance (ANOVA) test on the distance to the mean survival fraction for all the fluctuating communities against all the stable communities. This proved the statistical significance on the differences between the two groups, with the probability of observing these differences in a null model being very small ($p < 0.05$).

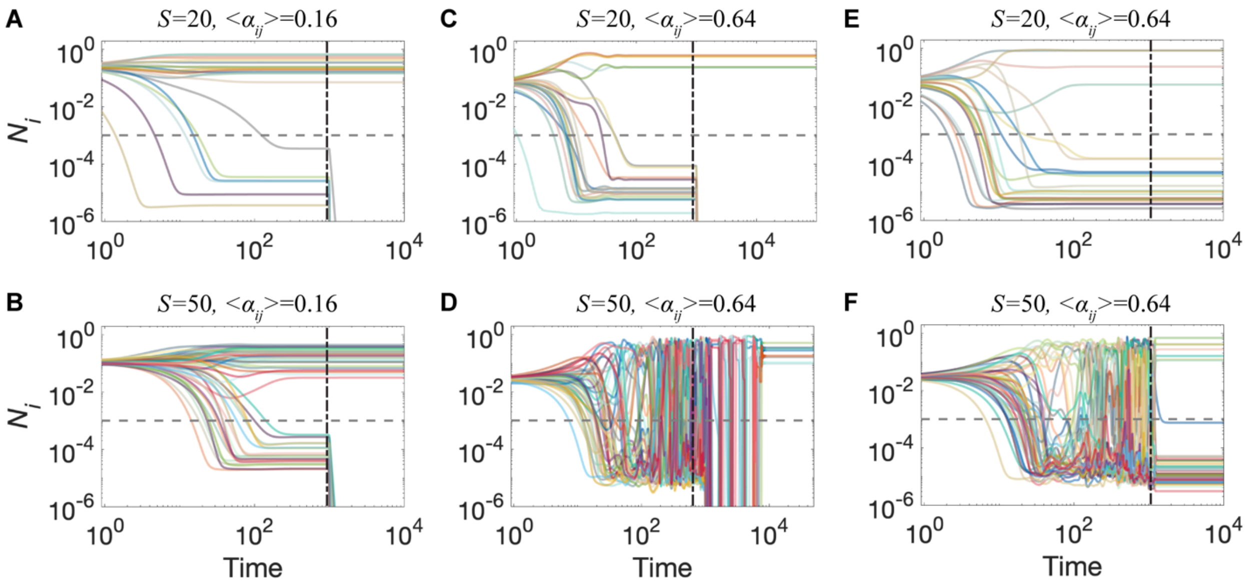


**Fig. S1. High diversity and persistent fluctuations allow and require each other, and are both sustained by dispersal.** (A)-(D) Representative time series for communities in which the dispersal rate is suddenly interrupted. At t=$10^3$ (vertical dashed line), the dispersal rate changes from D=$10^{-6}$ to D=0.0 for the rest of the simulation. (A) Before t=$10^3$, a community in phase I reaches a stable state with full coexistence. The dynamics after t=$10^3$ shows that interrupting dispersal does not significantly modify the abundances of the species. (B)-(C) Before t=$10^3$, communities in phase II reach an equilibrium in which species coexist at stable abundances, with some species laying below the extinction threshold. After stopping dispersal, only the species that are above the extinction threshold survive at stable abundances, and the rest undergo extinction. (D) A community in phase III exhibits persistent fluctuations while exposed to dispersal. After dispersal is interrupted, extinctions occur as species fall below the extinction threshold due to abundance fluctuations. After some time (approximately t=$10^4$) species extinctions have significantly reduced diversity in the community, and the surviving species reach a stable equilibrium. For the indicated parameter values, and over $10^3$ simulations, 90% of the simulated communities reached equilibrium after interrupting dispersal. (E-F) Representative time series for communities in which the most abundant species at t=$10^3$ is pinned (its abundance is artificially kept constant) for the rest of the simulation. (E) For communities that have reached stability, in this case in phase II, pinning the most abundant species has no effect on community dynamics. (F) In phase III, after a fast transient following the species pinning at t=$10^3$ (vertical dashed line), the community reaches a stable partial coexistence where some of the species lay below the extinction threshold. Out of $10^3$ simulations, 93% of the communities reached equilibrium after pinning the most abundant species.

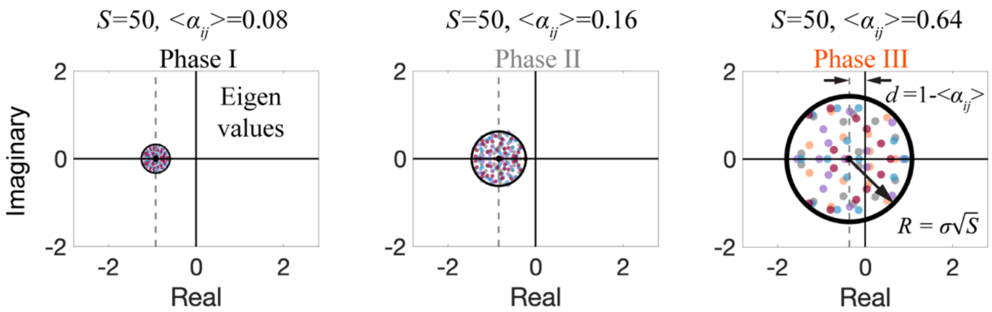


**Fig. S2. Unstable communities have (one or more) eigenvalues of the community matrix ($-\alpha_{ij}$) with positive real parts.** From left to right, the panels show the eigenvalues of representative community matrices for three different values of the average interaction strength. Within each panel, different colors correspond to the eigenvalues of 4 different community matrices. All the eigenvalues lie within a circle with radius $R$ centered at $d$ (*20*, *21*). For communities in phase III, where persistent fluctuations occur, some of the community matrix eigenvalues exhibit a positive real part. It was shown that the loss of stability of the equilibrium coincides with real parts of some community matrix ($-\alpha_{ij}$) eigenvalues becoming positive, although it is not the Jacobian matrix (*49*).

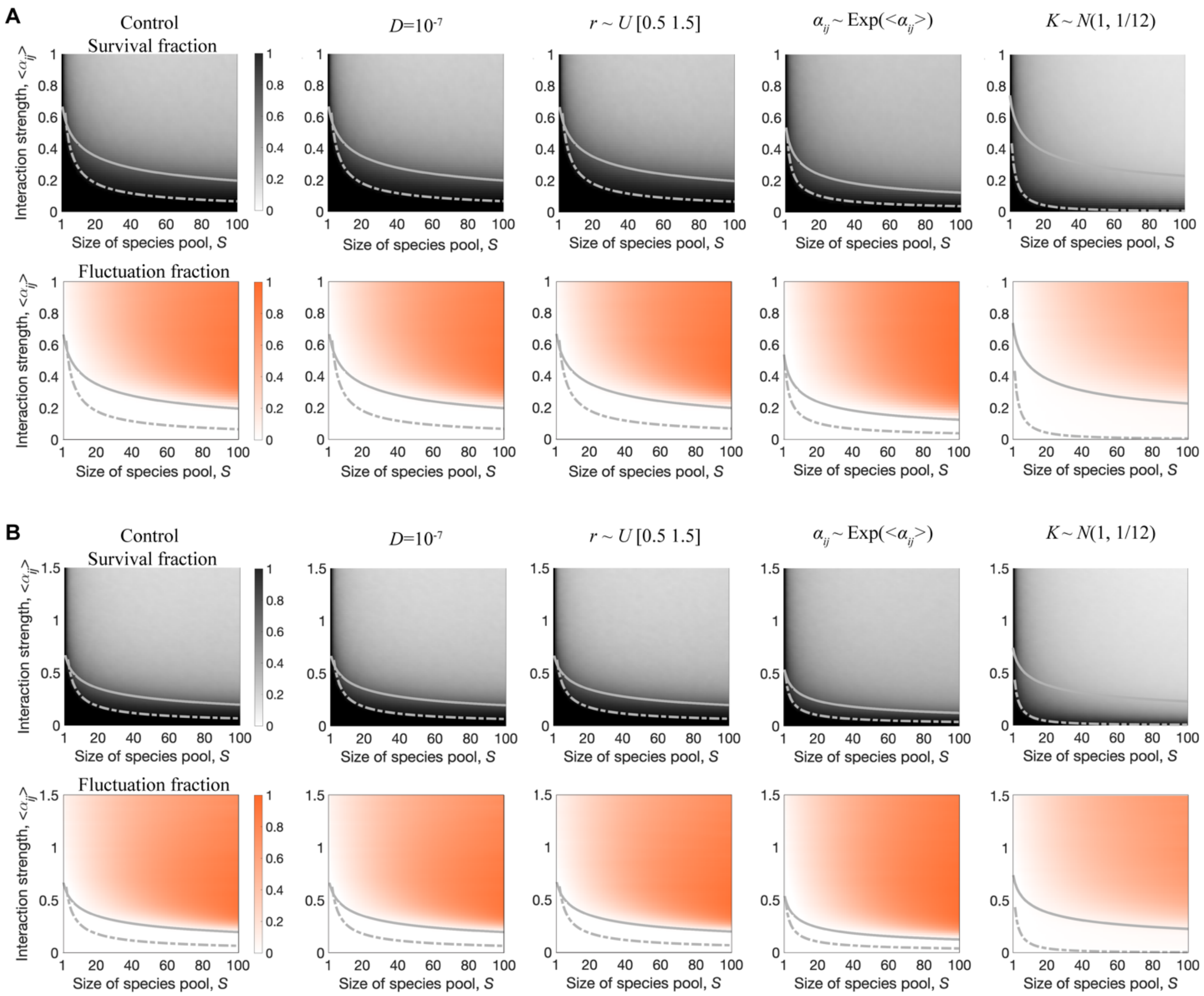


**Fig. S3. The three dynamical phases are robust to modeling choices.** (A) Panels on the left (Control) show the numerical (color map) and analytical (curves) phase diagrams as in Fig 1E and F. From left to right, the additional phase diagrams show the effects of lowering the dispersal rate to $D$=10$^{-7}$, sampling species growth rates from a uniform distribution, sampling interaction strengths from an exponential distribution, and sampling the carrying capacities from a Gaussian distribution. All non-specified parameter values are identical to the control case (Fig. 1E and F). (B) Phase diagrams analogous to those in (A), but for a higher average interaction strength <$\alpha_{ij}$ >= 1.5. Overall, these phase diagrams show that the three dynamical phases are qualitatively robust to different modeling choices (*27*).

k-Bacteria, p-Bacteroidota, c-Bacteroidia, o-Sphingobacteriales, f-Sphingobacteriaceae, g-Sphingobacterium
k-Bacteria, p-Firmicutes, c-Bacilli, o-Bacillales, f-Bacillaceae, g-Bacillus
k-Bacteria, p-Proteobacteria, c-Gammaproteobacteria, o-Burkholderiales, f-Oxalobacteraceae, g-Herbaspirillum
k-Bacteria, p-Proteobacteria, c-Gammaproteobacteria, o-Enterobacterales, f-Enterobacteriaceae, g-Raoultella
k-Bacteria, p-Proteobacteria, c-Gammaproteobacteria, o-Enterobacterales, f-Enterobacteriaceae, g-Pluralibacter
k-Bacteria, p-Proteobacteria, c-Gammaproteobacteria, o-Pseudomonadales, f-Pseudomonadaceae, g-Pseudomonas
k-Bacteria, p-Proteobacteria, c-Gammaproteobacteria, o-Enterobacterales, f-Yersiniaceae, g-Yersinia
k-Bacteria, p-Proteobacteria, c-Gammaproteobacteria, o-Xanthomonadales, f-Xanthomonadaceae, g-Stenotrophomonas
k-Bacteria, p-Proteobacteria, c-Gammaproteobacteria, o-Enterobacterales, f-Yersiniaceae
k-Bacteria, p-Proteobacteria, c-Gammaproteobacteria, o-Enterobacterales, f-Enterobacteriaceae, g-Citrobacter
k-Bacteria, p-Proteobacteria, c-Gammaproteobacteria, o-Xanthomonadales, f-Xanthomonadaceae, g-Stenotrophomonas
k-Bacteria, p-Firmicutes, c-Clostridia, o-Lachnospirales, f-Lachnospiraceae, g-Lachnospiraceae
k-Bacteria, p-Firmicutes, c-Bacilli, o-Lactobacillales, f-Streptococcaceae, g-Lactococcus
k-Bacteria, p-Proteobacteria, c-Gammaproteobacteria, o-Enterobacterales
k-Bacteria, p-Proteobacteria, c-Alphaproteobacteria, o-Rhizobiales, f-Rhizobiaceae, g-Allorhizobium
k-Bacteria, p-Proteobacteria, c-Gammaproteobacteria, o-Enterobacterales, f-Enterobacteriaceae
k-Bacteria, p-Firmicutes, c-Bacilli, o-Staphylococcales, f-Staphylococcaceae, g-Staphylococcus
k-Bacteria, p-Firmicutes, c-Bacilli, o-Staphylococcales, f-Staphylococcaceae, g-Staphylococcus
k-Bacteria, p-Firmicutes, c-Bacilli, o-Bacillales, f-Planococcaceae, g-Lysinibacillus
k-Bacteria, p-Proteobacteria, c-Alphaproteobacteria, o-Rhizobiales, f-Rhizobiaceae, g-Ochrobactrum
k-Bacteria, p-Firmicutes, c-Clostridia, o-Oscillospirales, f-Ruminococcaceae, g-Incertaesedis
k-Bacteria, p-Proteobacteria, c-Gammaproteobacteria, o-Enterobacterales, f-Erwiniaceae, g-Pantoea
k-Bacteria, p-Bacteroidota, c-Bacteroidia, o-Flavobacteriales, f-Weeksellaceae
k-Bacteria, p-Proteobacteria, c-Alphaproteobacteria, o-Acetobacterales, f-Acetobacteraceae, g-Roseomonas
k-Bacteria, p-Proteobacteria, c-Gammaproteobacteria, o-Alteromonadales, f-Alteromonadaceae, g-Alteromonas
k-Bacteria, p-Proteobacteria, c-Gammaproteobacteria, o-Oceanospirillales, f-Halomonadaceae, g-Cobetia
k-Bacteria, p-Proteobacteria, c-Gammaproteobacteria, o-Alteromonadales, f-Marinobacteraceae, g-Marinobacter
k-Bacteria, p-Proteobacteria, c-Alphaproteobacteria, o-Rhodobacterales, f-Rhodobacteraceae, g-Pacificibacter
k-Bacteria, p-Proteobacteria, c-Gammaproteobacteria, o-Enterobacterales, f-Erwiniaceae, g-Pantoea
k-Bacteria, p-Firmicutes, c-Bacilli, o-Staphylococcales, f-Staphylococcaceae
k-Bacteria, p-Proteobacteria, c-Gammaproteobacteria, o-Pseudomonadales, f-Moraxellaceae, g-Acinetobacter
k-Bacteria, p-Proteobacteria, c-Gammaproteobacteria, o-Enterobacterales, f-Enterobacteriaceae, g-Klebsiella
k-Bacteria, p-Proteobacteria, c-Gammaproteobacteria, o-Oceanospirillales, f-Nitrincolaceae, g-Neptunomonas
k-Bacteria, p-Bacteroidota, c-Bacteroidia, o-Flavobacteriales, f-Flavobacteriaceae, g-Maribacter
k-Bacteria, p-Bacteroidota, c-Bacteroidia, o-Flavobacteriales, f-Flavobacteriaceae, g-Winogradskyella
k-Bacteria, p-Proteobacteria, c-Gammaproteobacteria, o-Oceanospirillales, f-Halomonadaceae, g-Cobetia
k-Bacteria, p-Proteobacteria, c-Gammaproteobacteria, o-Enterobacterales, f-Enterobacteriaceae, g-Escherichia/Shigella
k-Bacteria, p-Proteobacteria, c-Gammaproteobacteria, o-Pseudomonadales, f-Pseudomonadaceae, g-Pseudomonas
k-Bacteria, p-Proteobacteria, c-Gammaproteobacteria, o-Enterobacterales
k-Bacteria, p-Proteobacteria, c-Gammaproteobacteria, o-Pseudomonadales, f-Pseudomonadaceae, g-Pseudomonas
k-Bacteria, p-Actinobacteriota, c-Actinobacteria, o-Micrococcales, f-Microbacteriaceae, g-Microbacterium
k-Bacteria, p-Firmicutes, c-Bacilli, o-Lactobacillales, f-Leuconostocaceae, g-Leuconostoc
k-Bacteria, p-Proteobacteria, c-Gammaproteobacteria, o-Xanthomonadales, f-Xanthomonadaceae, g-Stenotrophomonas
k-Bacteria, p-Proteobacteria, c-Gammaproteobacteria, o-Enterobacterales, f-Enterobacteriaceae, g-Klebsiella
k-Bacteria, p-Firmicutes, c-Bacilli, o-Lactobacillales, f-Streptococcaceae, g-Lactococcus
k-Bacteria, p-Proteobacteria, c-Gammaproteobacteria, o-Pseudomonadales, f-Pseudomonadaceae, g-Pseudomonas
k-Bacteria, p-Firmicutes, c-Bacilli, o-Brevibacillales, f-Brevibacillaceae, g-Brevibacillus
k-Bacteria, p-Firmicutes, c-Bacilli, o-Lactobacillales, f-Leuconostocaceae, g-Leuconostoc

**Fig. S4. Taxonomic identity of the 48 bacterial isolates.** The identities have been inferred from the ASV (Methods) of 16S samples taken from monocultures, which allow the classification of the 48 isolates down to the genus level. Colors are consistent with those in the main text and other supplementary figures.

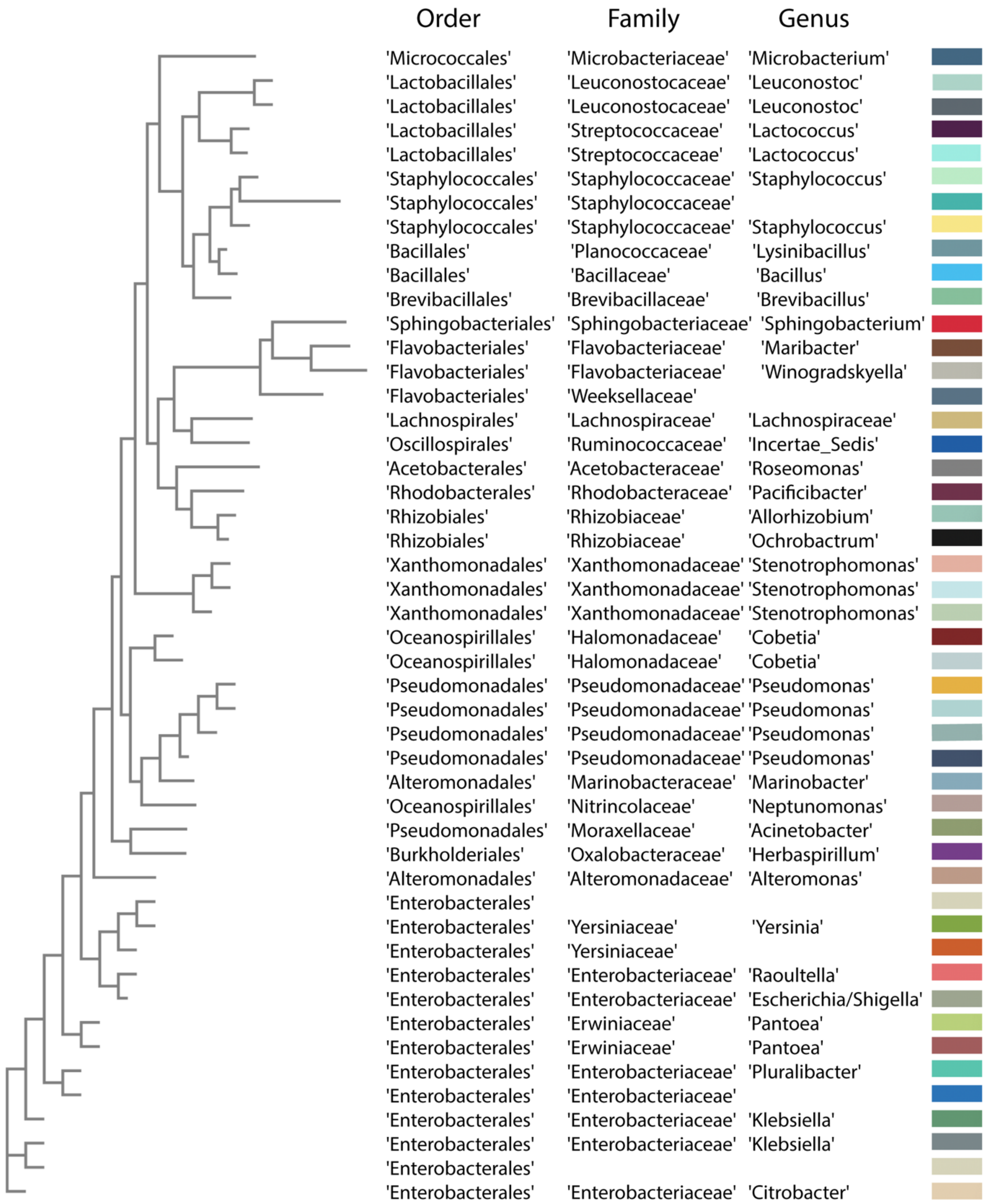


**Fig. S5. Phylogenetic tree of the 48 bacterial isolates.** The tree, generated with the Distance-matrix method from EMBL-EBL (*48*), shows the relative phylogenetic distance between the 48 bacterial isolates. The library contains bacterial isolates from either soil or *C. elegans* gut samples and spans 19 different orders and 26 different families. The rectangles display the color that is used in the figures (both in the Main Text and the SI) to show the abundance of each species in the different experiments.

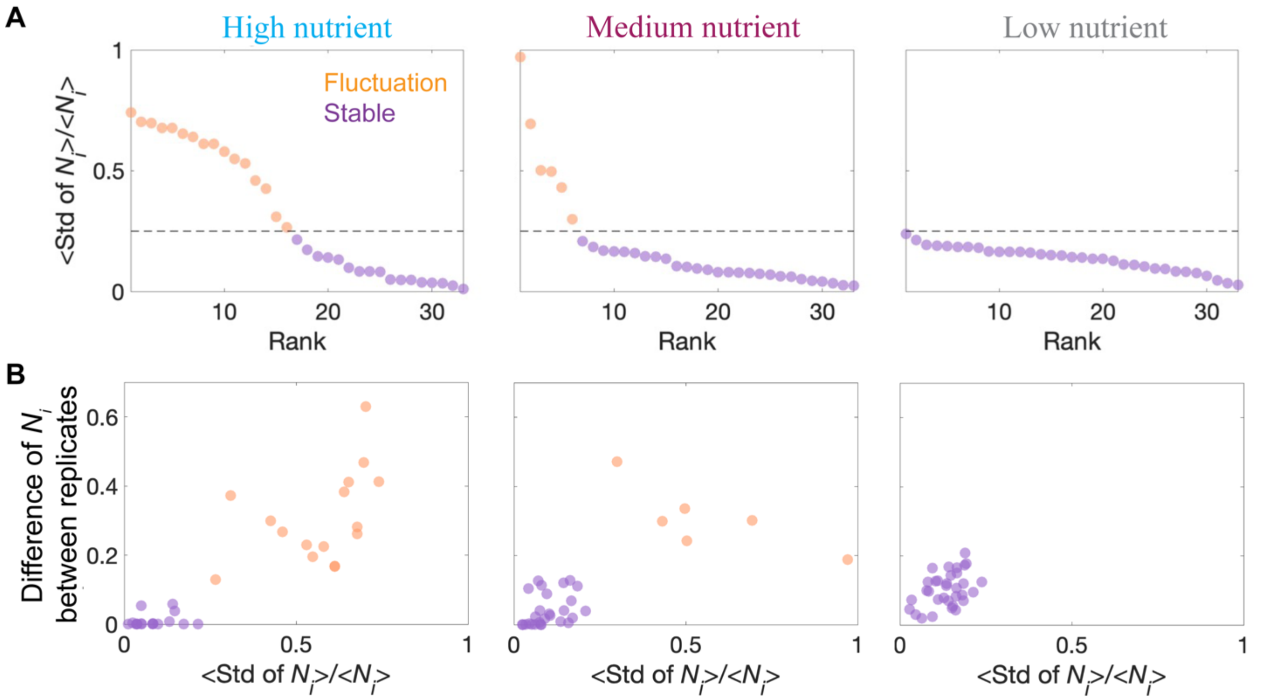


**Fig. S6. 2 different metrics consistently differentiate stable from fluctuating communities.** (A) The three panels show the coefficient of variation (see Methods) for species abundances in the experimental communities in the three different nutrients concentrations. We use a stability threshold of 0.25 (dashed line) to classify communities into stable (purple) and fluctuating (orange) ones. The number of fluctuating communities increases with the average interaction strength (nutrients concentration), with all the weakly interacting (low nutrients concentration) communities exhibiting stability. (B) Average difference (Euclidean distance) in the abundance of each species across replicate communities as a function of the community's coefficient of variation. Stable and fluctuating communities, defined as in (A), span in different regions, with stable communities clustering near the origin.

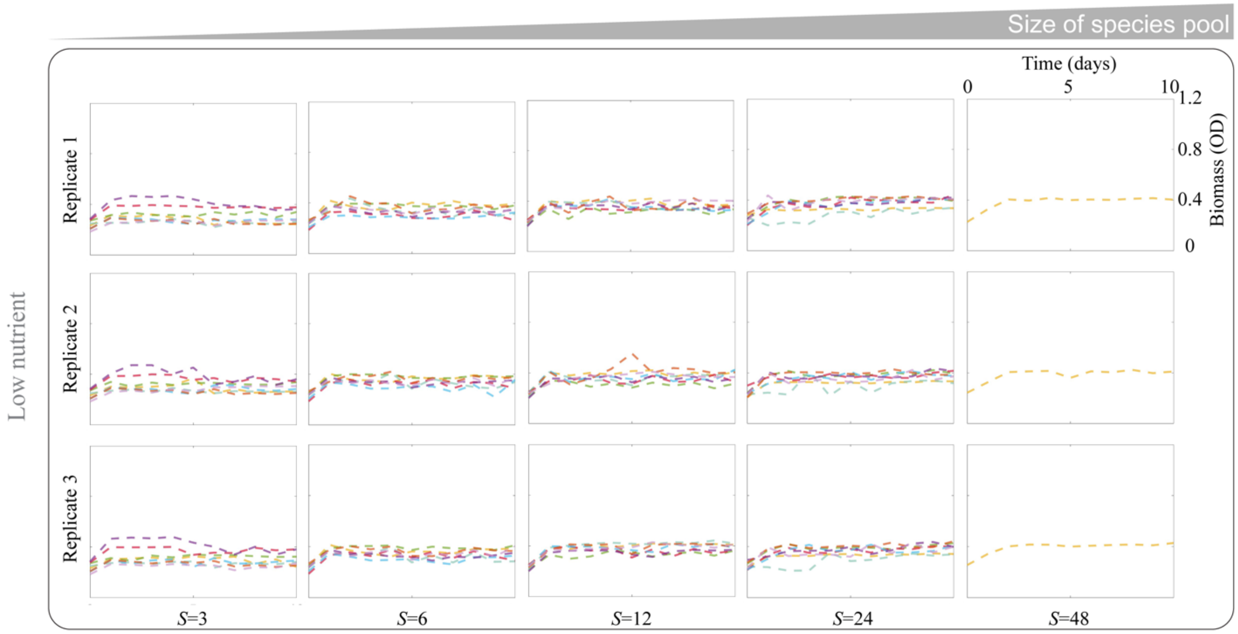


**Fig. S7. Total biomass reaches equilibrium in communities under low nutrients concentration (low interaction strength)**. Each panel shows the time series for the OD (600nm) of the eight communities with different species pool composition (depicted by different colors). Each column stands for a different species pool size S (for the case of S=48, there is only one community containing the full library of bacterial species). Each row shows the data for a different replicate of the experiment.

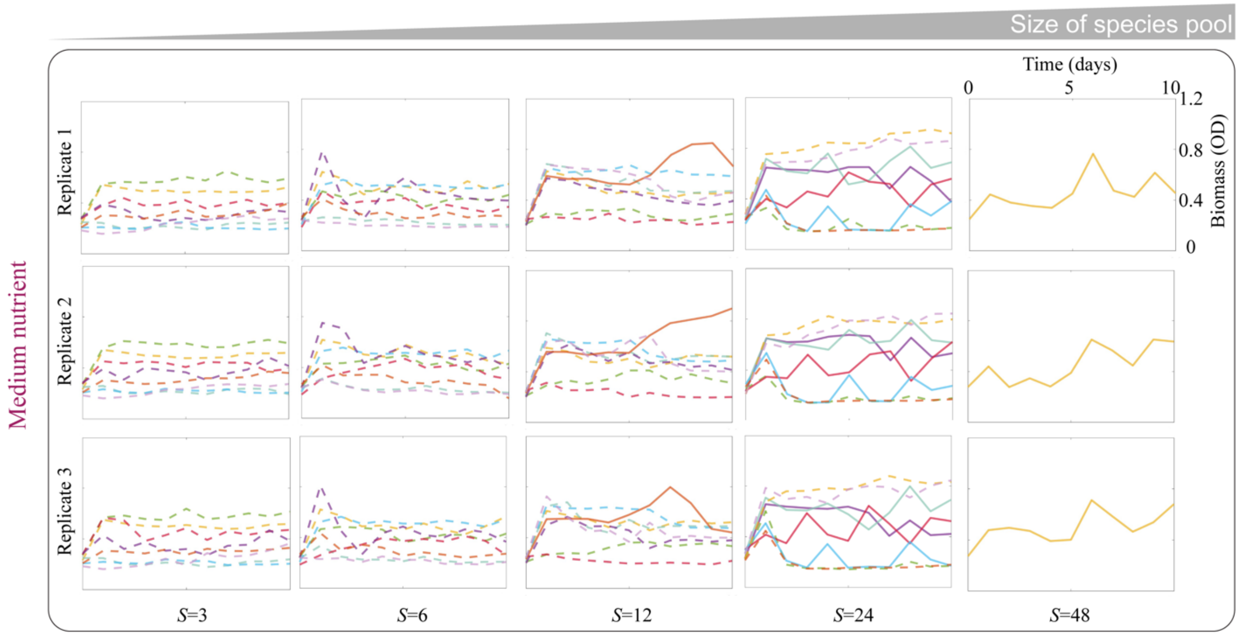


**Fig. S8. Increasing the species pool size leads to persistent fluctuations in total biomass under intermediate nutrients concentration (intermediate interaction strength).** Each panel shows the time series for the OD (600nm) of the eight communities with different species pool composition (depicted by different colors). Each column stands for a different species pool size S (for the case of S=48, there is only one community containing the full library of bacterial species). Each row shows the data for a different replicate of the experiment. Solid lines (dashed lines) represent fluctuating (stable) communities, the OD fluctuations between day 7 and day 10 were considered to differentiate fluctuating and stable communities as shown in Fig. S6.

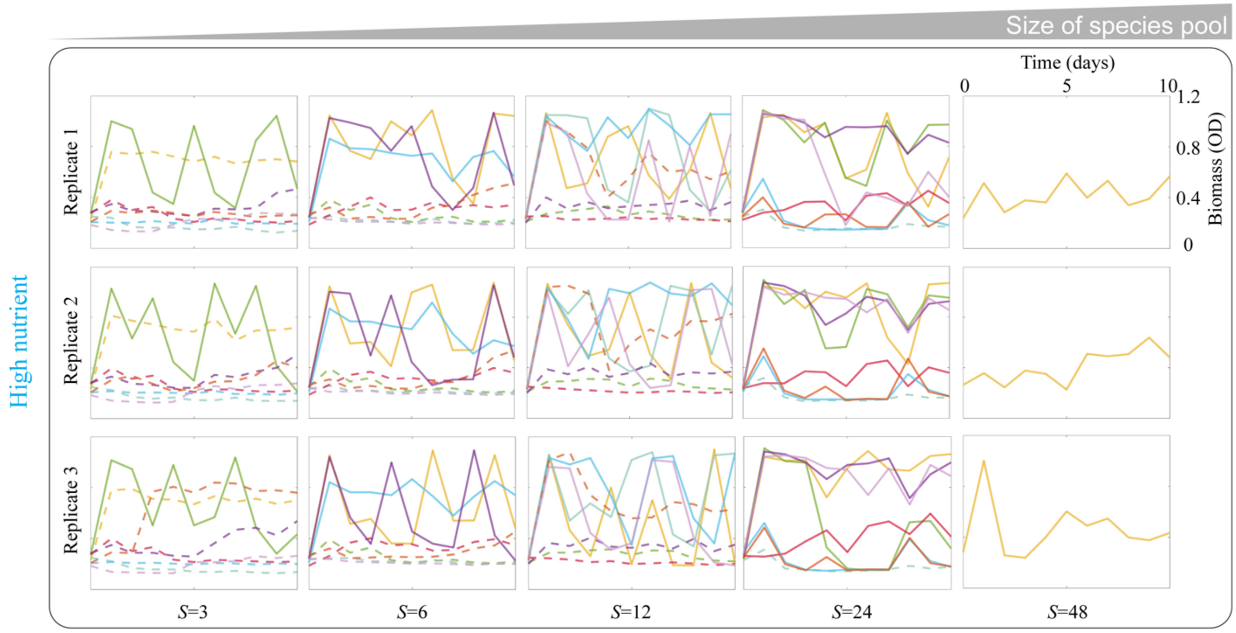


**Fig. S9. Increasing the species pool size leads to persistent fluctuations in total biomass under high nutrients concentration (high interaction strength).** Each panel shows the time series for the OD (600nm) of the eight communities with different species pool composition (depicted by different colors). Each column stands for a different species pool size S (for the case of S=48, there is only one community containing the full library of bacterial species). Each row shows the data for a different replicate of the experiment. Solid lines (dashed lines) represent fluctuating (stable) communities, the OD fluctuations between day 7 and day 10 were considered to differentiate fluctuating and stable communities as shown in Fig. S6.

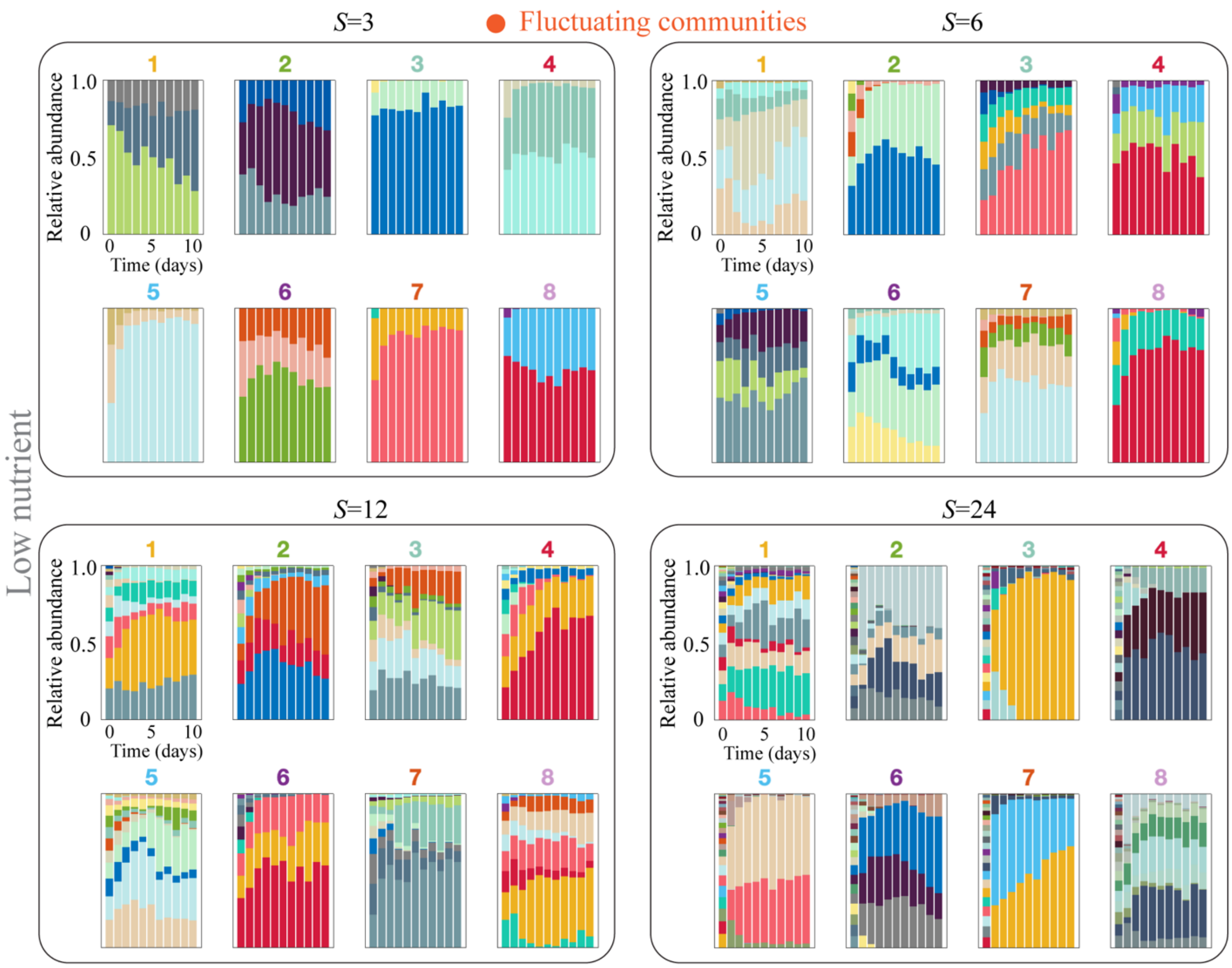


**Fig. S10. Time series for the relative species abundances of the experimental communities with low average interaction strength (low nutrients concentration).** Each panel shows the full time series for each of the 8 communities with the indicated species pool size (S=3, 6, 12 and 24). Bar colors stand for species identities as in Fig. S4. Under this nutrients condition, all of the communities reached a stable equilibrium (Methods). The color of the number on the top of each panel corresponds to the color assigned to the same community in Fig. S7.

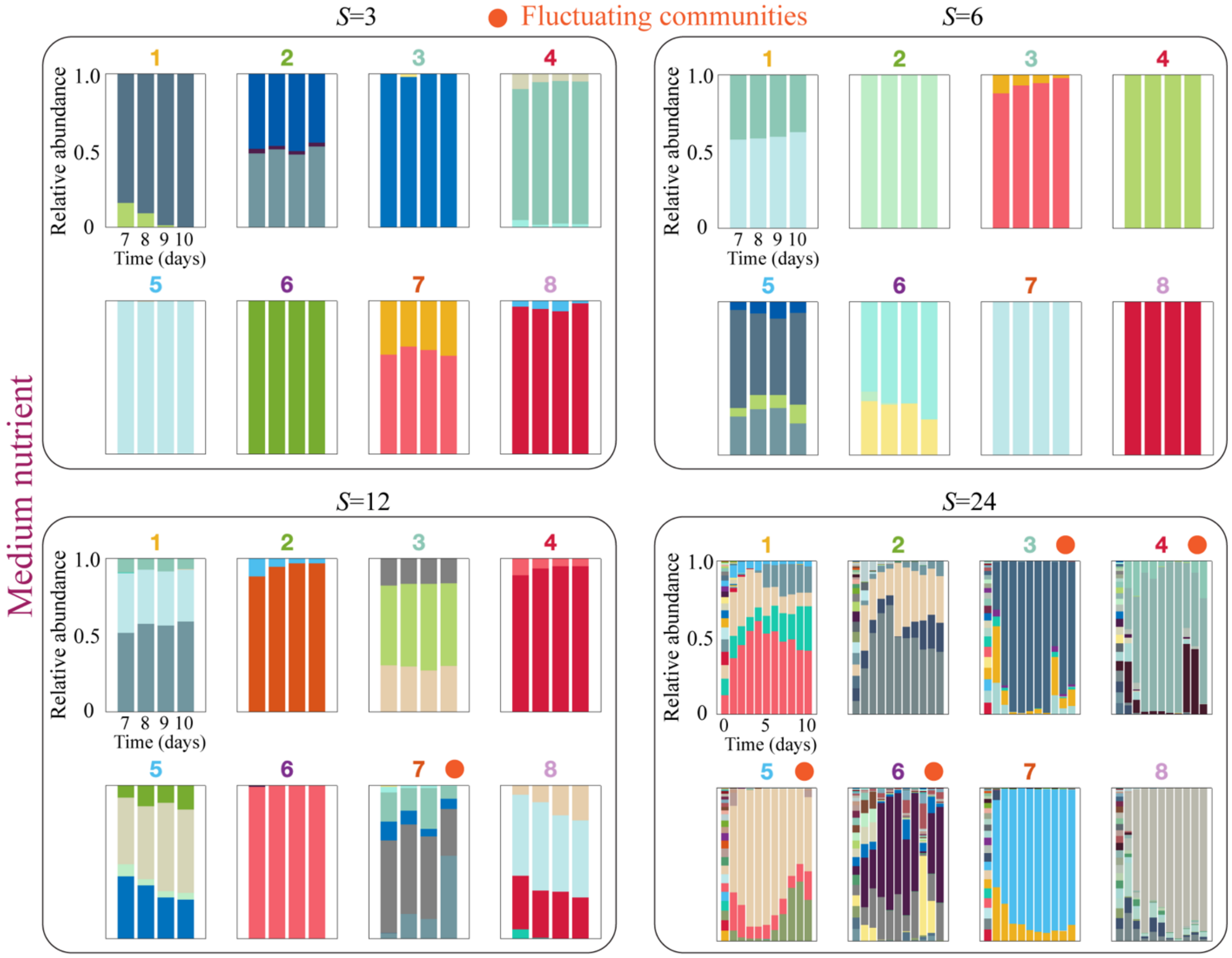


**Fig. S11. Time series for the relative species abundances of the experimental communities with intermediate average interaction strength (medium nutrients concentration).** Each panel shows the full time series for each of the 8 communities with the indicated species pool size (S=3, 6, 12 and 24). Bar colors stand for species identities. The orange dot on top of some panels indicates that the community exhibits persistent fluctuations (Methods). The color of the number on the top of each panel corresponds to the color assigned to the same community in Fig. S8. For S= 3, 6, and 12, we only sequenced samples of the last 4 days (7 to 10) of the experiment.

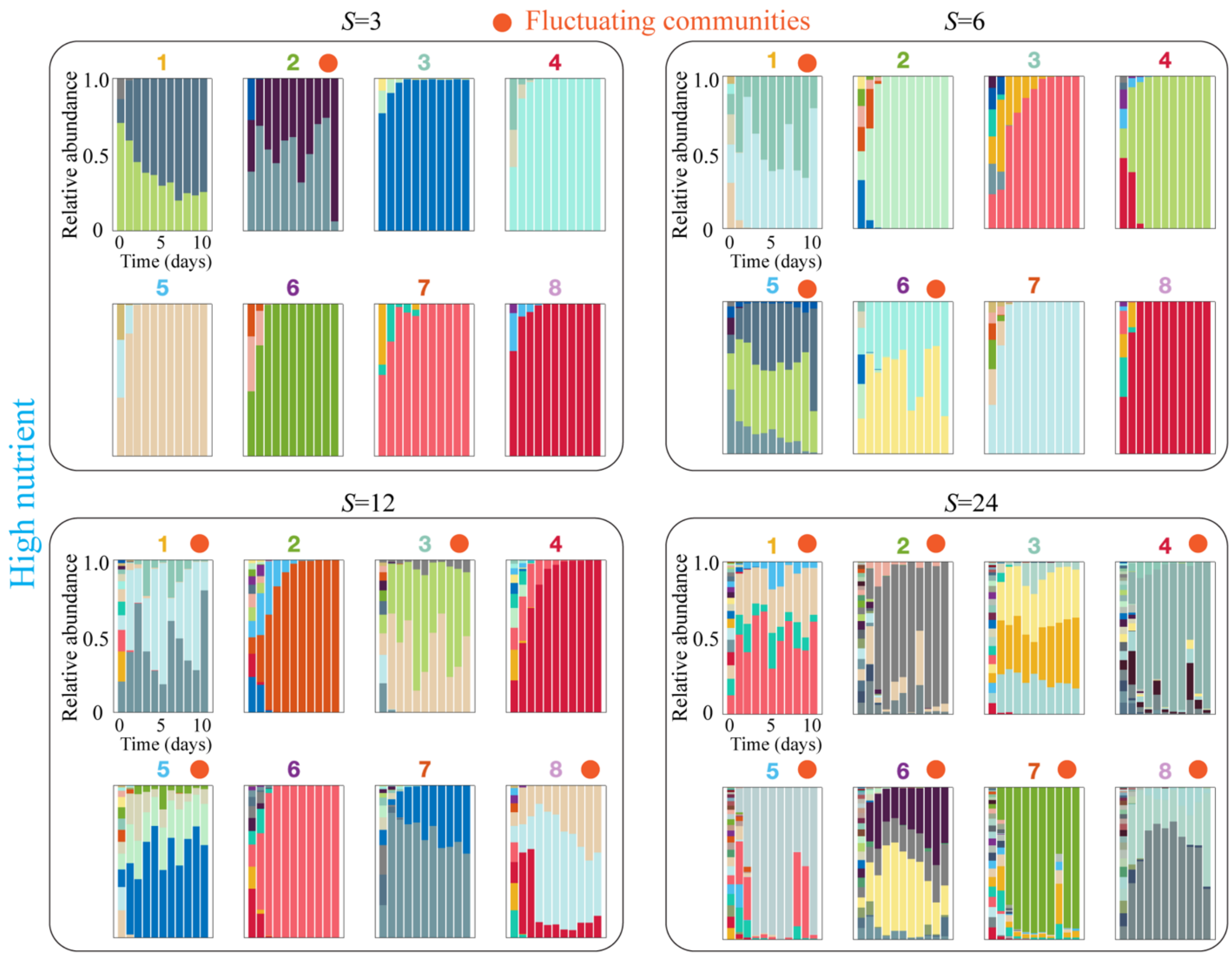


**Fig. S12. Time series for the relative species abundances of the experimental communities with high average interaction strength (high nutrients concentration).** Each panel shows the full time series for each of the 8 communities with the indicated species pool size (S=3, 6, 12 and 24). Bar colors stand for species identities. The orange dot on top of some panels indicates that the community exhibits persistent fluctuations (Methods). The color of the number on the top of each panel corresponds to the color assigned to the same community in Fig. S9.

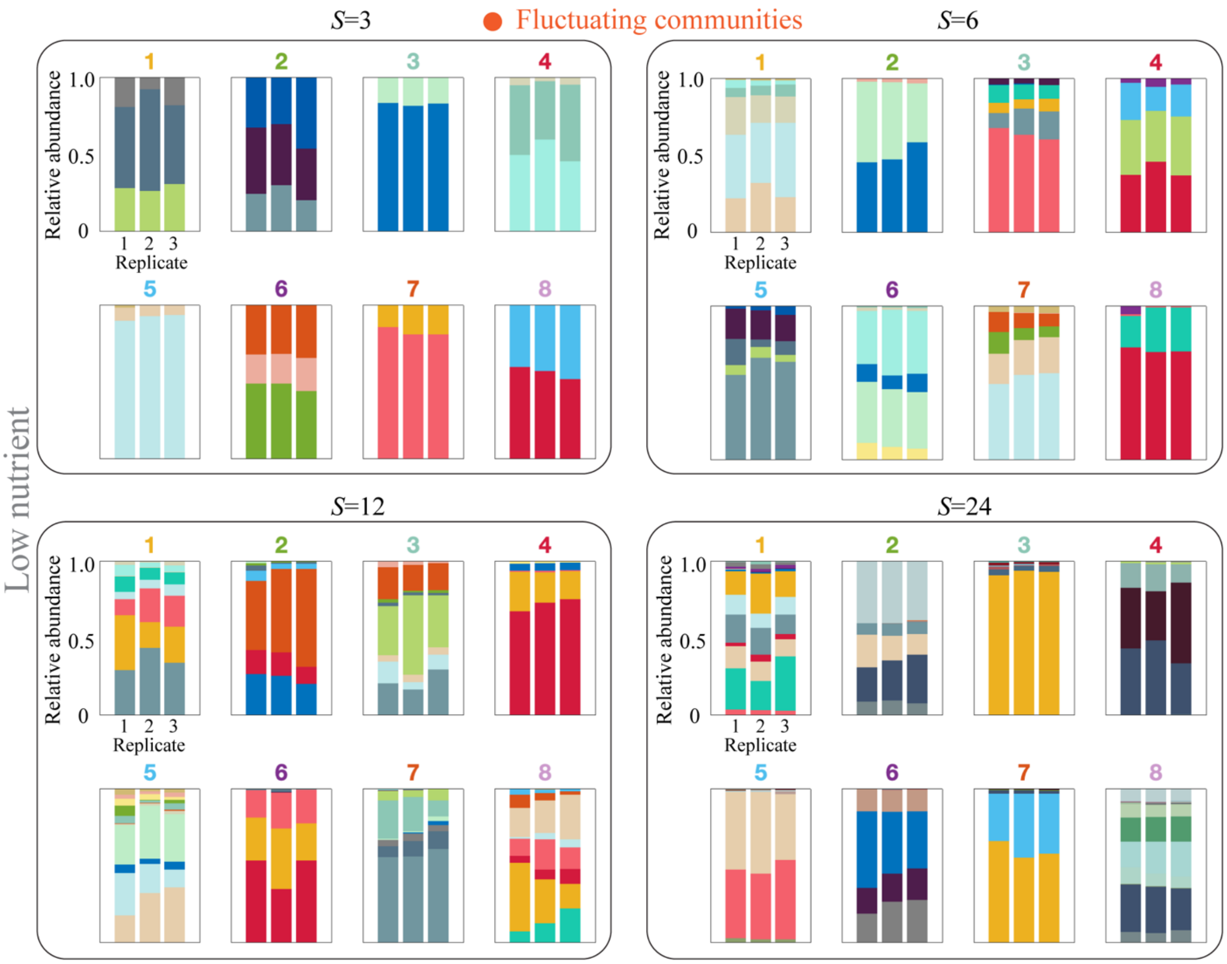


**Fig. S13. Species abundance at the end of the experiment under low nutrients concentration.** Each panel shows the relative species abundances at the end experiment for each of the 3 replicate communities across 8 different compositions of the species pool for each species pool size (S=3, 6, 12 and 24). Bar colors stand for species identities. The orange dot on top of some panels indicates that the community exhibits persistent fluctuations (Methods). The color of the number on the top of each panel corresponds to the color assigned to the same community in Fig. S7.

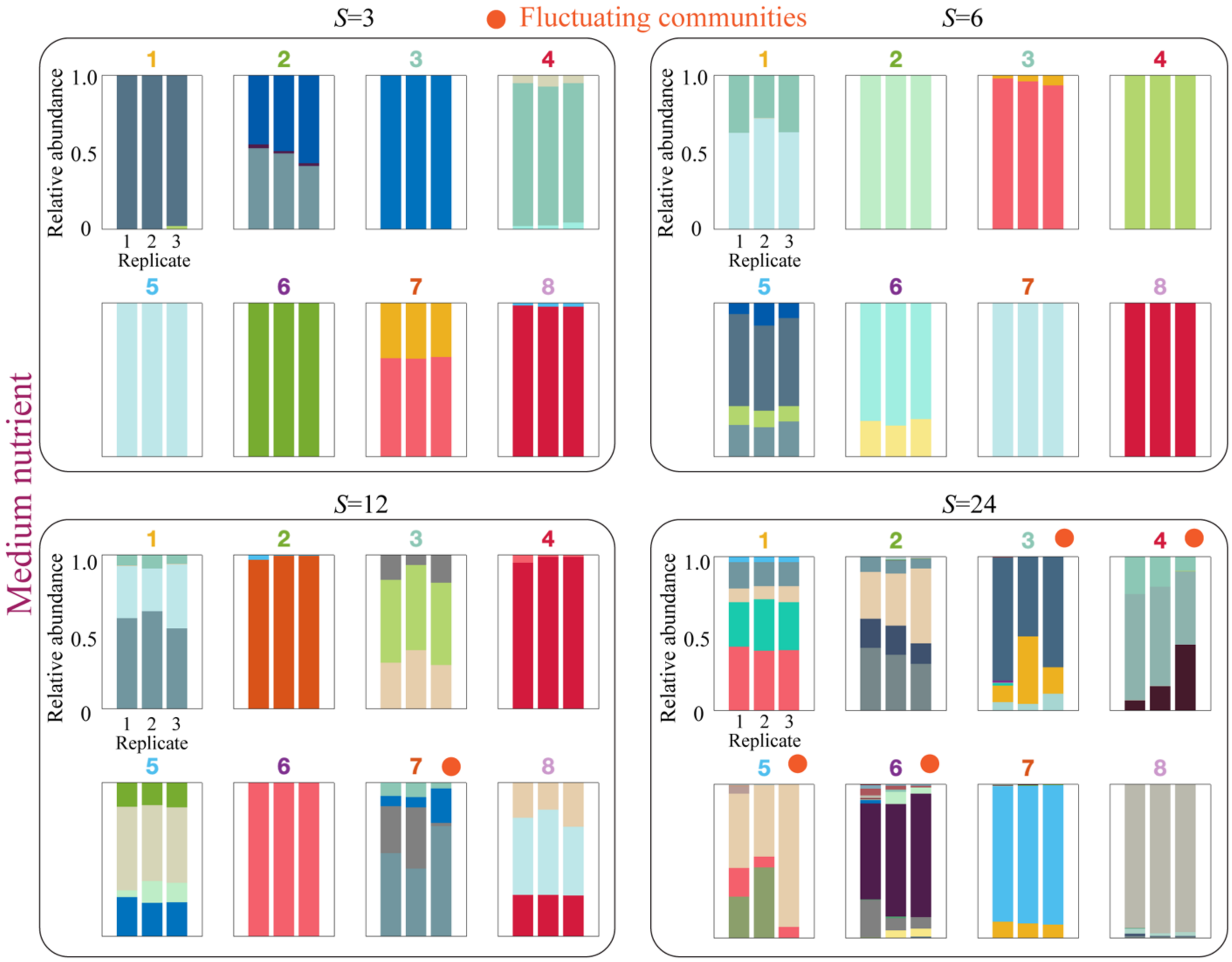


**Fig. S14. Species abundance at the end of the experiment under medium nutrients concentration.** Each panel shows the species relative abundances at the end experiment for each of the 3 replicate communities across 8 different compositions of the species pool for each species pool size (S=3, 6, 12 and 24). Bar colors stand for different species identities. The orange dot on top of some panels indicates that the community exhibits persistent fluctuations (Methods). The color of the number on the top of each panel corresponds to the color assigned to the same community in Fig. S8.

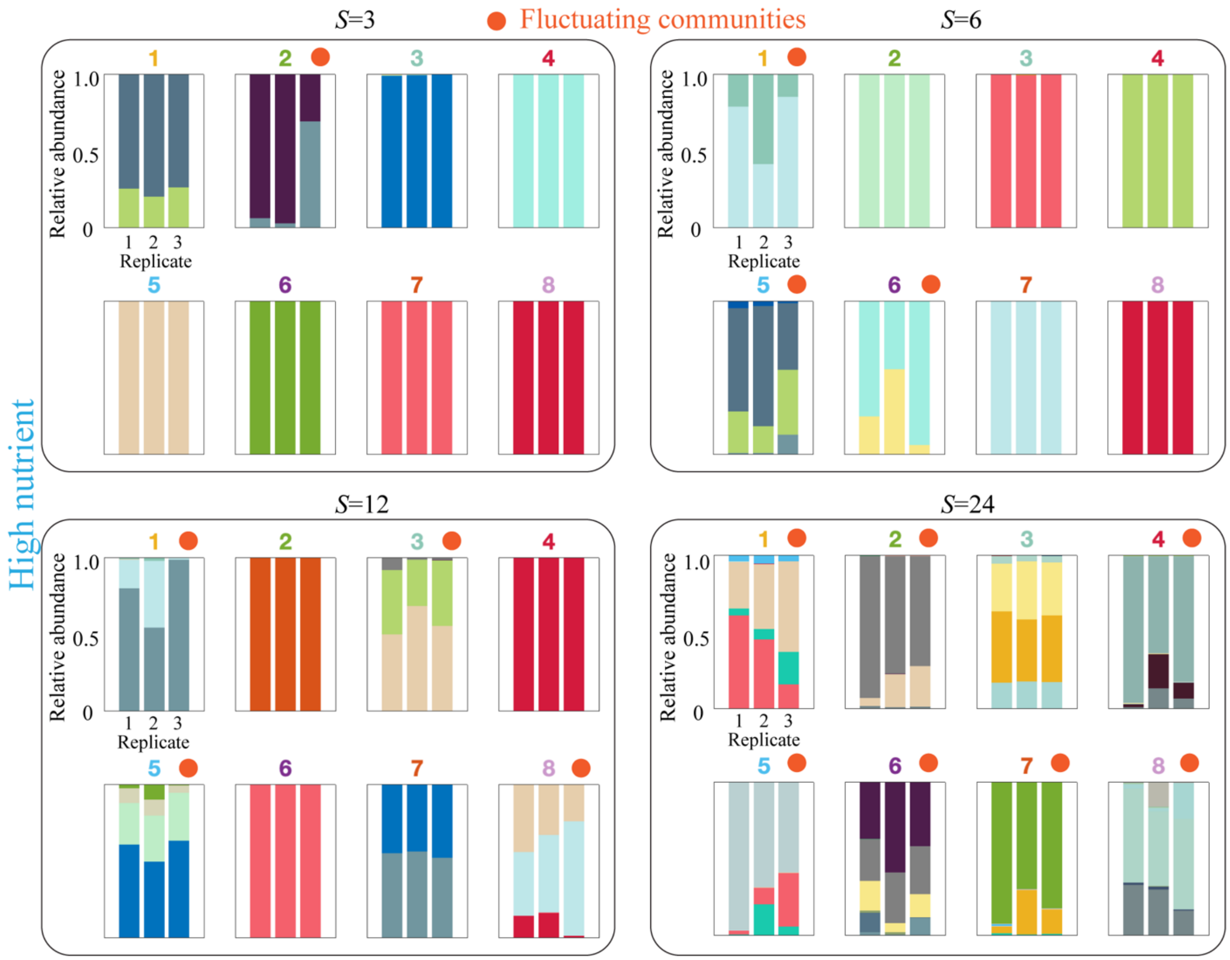


**Fig. S15. Species abundance at the end of the experiment under high nutrients concentration.** Each panel shows the species relative abundances at the end experiment for each of the 3 replicate communities across 8 different compositions of the species pool for each species pool size (S=3, 6, 12 and 24). Bar colors stand for different species identities. The orange dot on top of some panels indicates that the community exhibits persistent fluctuations (Methods). The color of number on the top of each panel corresponds to the color assigned to the same community in Fig. S9.

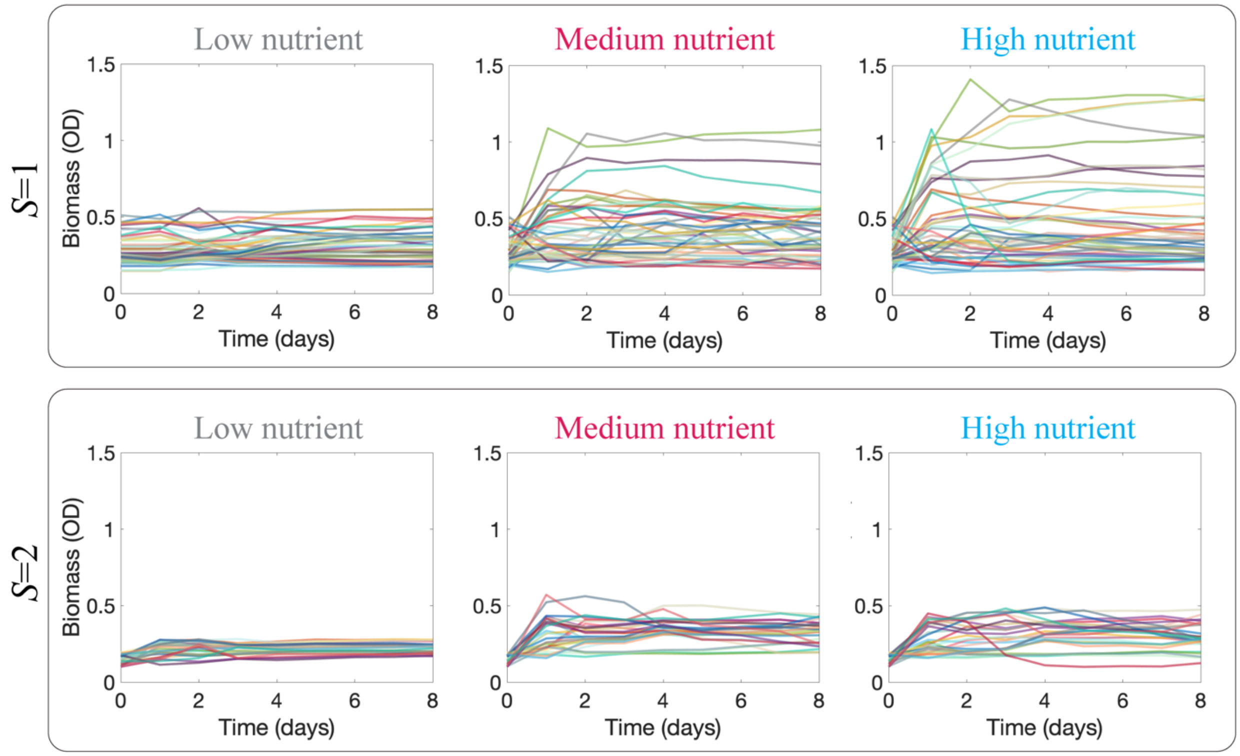


**Fig. S16. Monocultures and 2-species cocultures tend to reach stability in total biomass.** On top, monocultures tend to reach a stable OD (600nm) value at the end of each daily cycle. The width of the observed range of OD values increases with nutrient concentration (low, medium, and high, from left to right). On bottom, time series for the OD (600nm) of 30 different species pair cocultures. The variability of the OD reached in pairwise coculture also increases with nutrient concentration, but to a less extent than it does for monocultures. Different colors stand for different species identities (top) and different species pairs (bottom).

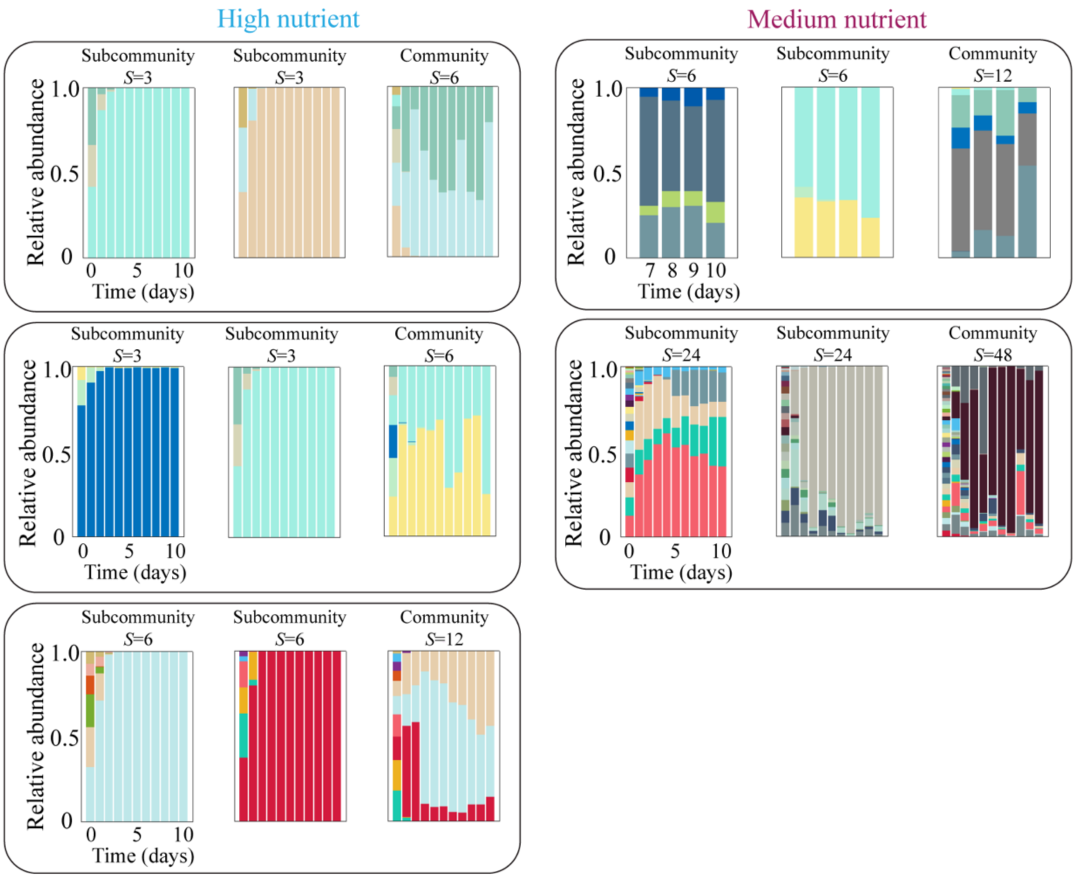


**Fig. S17. Increasing species pool size can lead to emergent fluctuations in species abundances.** The panels show representative examples in which a pair of different communities reach stability, while a community with larger species pool, composed by all the species present in that pair, exhibits persistent fluctuations. Each rectangle encloses a different example involving a specific set of communities and experimental condition (nutrients concentration). The top right rectangle shows data for the last 4 days of the experiment (the only days in which these communities were sampled for sequencing), and the rest show the full time series for the 10-day experiment.

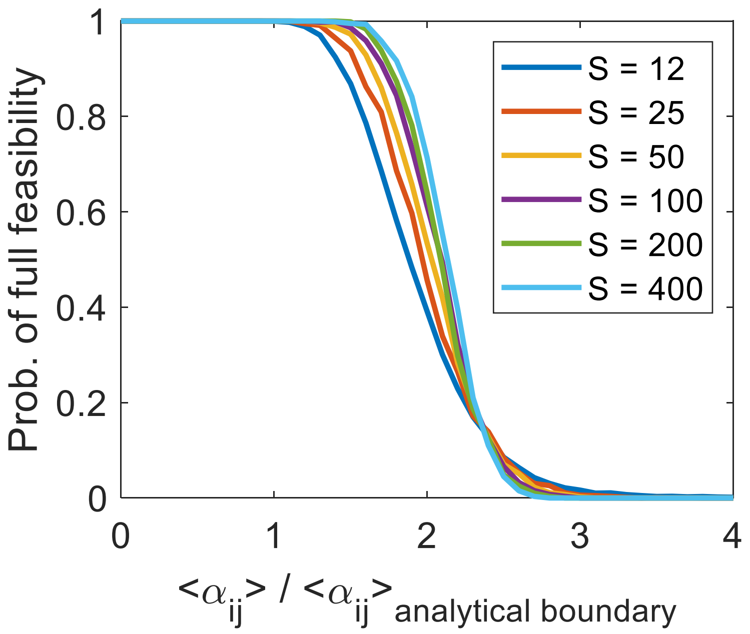


**Fig. S18. The probability of full coexistence as a function of the mean interaction strength $\langle \boldsymbol{\alpha_{ij}} \rangle$ exhibits a sharp phase transition between phases (I) and (II) when $\boldsymbol{S}$ is large in simulations.** The x-axis is normalized by $\langle \alpha_{ij} \rangle$ where the analytical survival boundary is expected. While all curves decrease to zero in the same region, the width of the crossover regime becomes narrower with increasing $S$. The fact that all curves decrease to zero at the same region, shows that the analytical expression indeed captures the correct dependence of the boundary in $\langle \alpha_{ij} \rangle$ on $S$.
